# Telomere-to-Telomere Accurate and Gapless Korean Standard Reference Genome

**DOI:** 10.1101/2025.11.17.688257

**Authors:** Jihun Bhak, Yoonsung Kwon, Dong-Hyun Shin, Kyungwhan An, Hyoungjin Choi, Sangsoo Park, Changhan Yoon, Yookyung Choi, Hyomin Lee, Daeui Park, Sungwon Jeon, Yeonsu Jeon, Jongbum Jeon, Soobok Joe, Jin Ok Yang, Jong-Hwan Kim, Yun Sung Cho, Jungeun Kim, Kyun Shik Chae, Chang Geun Kim, Bayarsaikhan Delger, Jaesuk Lee, Semin Lee, Haeyoung Jeong, Jong Bhak

## Abstract

We present KOREF1-G-TTAGGA, the first Telomere-to-Telomere Accurate and Gapless Genome Assembly, standing as the Korean standard reference genome. The paternal and maternal haplotypes spanned 2.91 and 3.03 Gb. Genome-wide, at least 99.15% of assembled sequences were reliably haplotype-resolved, and over 95% remained accurate even in the most error-prone loci, including centromeric satellite arrays and segmental duplications. Rare-k-mer copy-number concordance within satellite arrays held up 98.9% and 98.8% per haplotype. Bionano optical maps fully spanned all canonical rDNA arrays on the five acrocentric chromosomes. The assembly quality index (AQI) of 99.77 and 99.69 exceeded the reference-quality threshold of 90, and more than 99.99% of gene and cCRE sequence was free of structural error. Both haplotypes further showed high base-level accuracy, with consensus quality value (QV) of 81.19 and 79.03, corresponding to one error per 131 and 80 Mb. KOREF1-G-TTAGGA is among the highest-quality East Asian telomere-to-telomere assemblies. It moreover anchors a decade-spanning multi-ome reference dataset for defining individual’s molecular states. Together, these resources define the personal referenceome as a foundation for individual biology and precision medicine, with the assemblies, annotations, and all multi-omic data openly available at https://koreanreference.org.

## Background & Summary

Understanding an individual’s biology begins with a personal genome^1,2^. Yet personal genome sequences have long been represented only indirectly, as a list of variants called against the human standard reference genomes^3–12^. Those pseudo-haploid references, GRCh38 and T2T-CHM13, derived from multiple donors and a hydatidiform mole cell line, respectively, incompletely represent the diploid genome of a human^13–20^. They cannot capture the full spectrum of genetic variants in diverse individuals, especially in underrepresented populations^21–27^. Population-specific reference genomes were therefore developed to improve read mapping and variant discovery for individuals in such groups^22–24,28,29^. More recently, pangenome references have pushed further, enabling more comprehensive genotyping even at structurally complex loci^30–36^. However, pangenome references still cannot reconstruct the complete genome sequence of any one person^34,35,37^. Even the most comprehensive personal multi-omic studies have been built on a human standard reference genome rather than on the individual’s own genome^38–43^. Personal genomics, for all its name, has yet to be practiced on a complete personal reference genome.

Each person’s own genome is the standard reference for their own biology. The Personal Genome Project (PGP), launched by George Church in 2005, established the personal genome as a foundation for interpreting one individual’s biology^1^. The KOrean REFerence (KOREF) project carried the principle into a national effort in 2006, governed by the Ministry of Science and ICT (MSIT) and Korea Bioinformation Center (KOBIC), setting out to build the Korean standard reference genome and multiome^22,44,45^. Its founding sample, KOREF1 (PGP hu3D760A; KPGP KPGP9), anchored a single-individual Korean reference (KOREF-S). A Korean pan-genome consensus (KOREF-C), built on KOREF-S from 40 Korean variomes, has become the standard reference of Korean genomic diversity. It mapped variation within Koreans, identified Korean-specific sequences, and improved variant detection, taking a first step toward personal genomics for every Korean^22^. Yet neither reference was a complete sequence, and even the trio-phased assembly of 2022 remained unresolved regions^22,45^.

A complete genome alone is not enough to study an individual’s biology. Personal multi-omics must be measured against that individual’s own genome^42^. Mapped to a personal diploid genome instead of the global standard reference genome, an individual’s data reveals more than a million allele-specific loci that the reference misses^42^. No computational model yet predicts these states at the level of an individual^46–48^. Sequence-to-function deep learning and state-of-the-art AI models are trained on the reference genome, so they predict population averages rather than individuals^46–51^. Across individuals, their predictions correlate poorly with measured expression, and they often get the direction of a variant’s effect wrong^46–48^. Personal biology therefore requires both a personal reference genome and its own multi-omic reference dataset that defines the individual’s molecular states and variations^52^. Testing that biology also requires experimentally accessible cells that a living donor cannot repeatably provide^53^. Renewable iPSCs and cell lines from the same individual supply exactly that^54^.

Here, we present KOREF1-G-TTAGGA, a Telomere-to-Telomere Accurate and Gapless Genome Assembly (TTAGGA) of KOREF1. It is the Korean standard reference genome integrated with a decade-spanning multiome (Fig. 1 and Table 1). It also comes with reference materials from the same individual, namely a B lymphoblastoid cell line (Korean Cell Line Bank; KCLB #60211) and an induced pluripotent stem cell (iPSC) line. The KOREF project follows the open model of the PGP, so its genome, multiomic data, and cell lines are released without any restriction (https://koreanreference.org), as a shared resource for the community^1,6,40^. Together, these resources define the personal referenceome as a foundation for studying individual biology and, in turn, for precision medicine.

**Figure 1.**
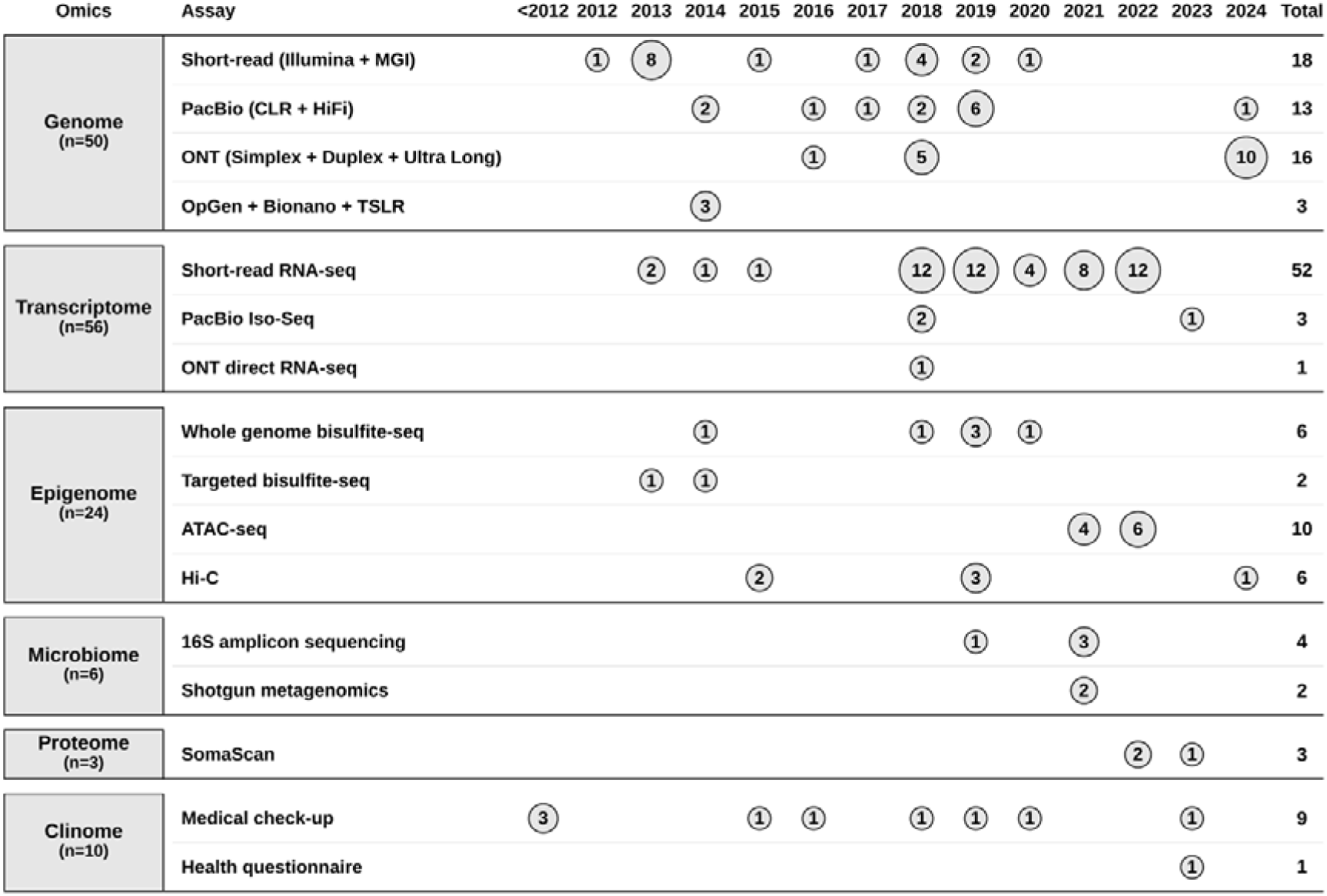
Over thirteen years of longitudinal multi-omic data acquisition from a single individual. Each circle marks the year an assay was performed, with the enclosed number giving the number of experiments generated that year; row totals are given at right and layer totals in the left margin. Data span more than a decade of repeated sampling of the same individual across genome (n = 50), transcriptome (n = 56), epigenome (n = 24), microbiome (n = 6), proteome (n = 3), and clinome (n = 10). Successive sequencing platforms enter the record as they became available, short read from 2012, PacBio from 2014, ONT from 2016, with ultra-long ONT concentrated in 2024, while transcriptome and epigenome assays were added from 2013 onward, yielding a cumulative multi-omic resource anchored to a single genome.

**Table 1.** Multiomic datasets generated for the KOREF1 individual.

| Omics | Assay | Biosource | Experiments | Yield (bases) | Platform |
| --- | --- | --- | --- | --- | --- |
| Genome | Short-read (Illumina + MGI) | Blood; LCL | 18 | 2.55 Tb | Illumina: HiSeq 2000, HiSeq 4000, NovaSeq 6000; MGI: BGISEQ-500, DNBSEQ-G400, DNBSEQ-T7 |
|  | PacBio (CLR + HiFi) | Blood | 13 | 2.32 Tb | PacBio: RS II, Sequel, Sequel II, Revio |
|  | ONT (Simplex + Duplex + Ultra Long) | Blood; LCL | 16 | 1.66 Tb | ONT: MinION, PromethION |
|  | OpGen + Bionano + TSLR | Blood | 3 | 16.3 Gb | OpGen: OpticalMapping; BioNano: Nanochannel; Illumina: HiSeq 2500 |
|  |  |  | **50** | **6.55 Tb** |  |
| Transcriptome | Short-read RNA-seq | Blood; LCL; Platelet | 52 | 624 Gb | Illumina: HiSeq 2500, NovaSeq 6000; MGI: BGISEQ-500, DNBSEQ-T7 |
|  | PacBio Iso-Seq | Blood; LCL | 3 | 69.9 Gb | PacBio: Sequel, Sequel II |
|  | ONT direct RNA-seq | LCL | 1 | 440 Mb | ONT: MinION |
|  |  |  | **56** | **695 Gb** |  |
| Epigenome | Whole genome bisulfite-seq | Blood | 6 | 700 Gb | Illumina: HiSeq 2000, HiSeq X, NovaSeq 6000 |
|  | Targeted bisulfite-seq | Blood | 2 | 25.3 Gb | Illumina: HiSeq 2000 |
|  | ATAC-seq | PBMC | 10 | 198 Gb | Illumina: NovaSeq 6000 |
|  | Hi-C | LCL; PBMC | 6 | 1.31 Tb | Illumina: HiSeq X, NovaSeq 6000, NovaSeq X Plus |
|  |  |  | **24** | **2.23 Tb** |  |
| Microbiome | 16S amplicon sequencing | Gut; Oral | 4 | 520 Mb | Illumina: MiSeq |
|  | Shotgun metagenomics | Oral | 2 | 20.3 Gb | Illumina: NovaSeq 6000; MGI: DNBSEQ-G400 |
|  |  |  | **6** | **20.8 Gb** |  |
| Proteome | SomaScan | Plasma | 3 | — | SomaLogic: SomaScan |
|  |  |  | **3** | **—** |  |
| Clinome | Medical check-up | — | 9 | — | Medical Check-up |
|  | Health questionnaire | — | 1 | — | Health Questionnaire |
|  |  |  | **10** | **—** |  |
| Sequencing total |  |  | **136** | **9.50 Tb** |  |
| All experiments |  |  | **149** | **—** |  |

Experiments and assays across six omics layers, all from one individual. For each assay the table lists the biosource, the number of experiments, the total sequence yield in bases, and the platforms used. Genome data comprise short-read, PacBio (CLR and HiFi), ONT, and the OpGen, BioNano and synthetic long-read platforms (50 experiments, 6.55 Tb); transcriptome data comprise short-read RNA-seq, PacBio Iso-Seq and ONT direct RNA-seq (56 experiments, 695 Gb); epigenome data comprise whole-genome and targeted bisulfite sequencing, ATAC-seq and Hi-C (24 experiments, 2.23 Tb); microbiome data comprise 16S amplicon sequencing of gut and oral samples and shotgun metagenomic sequencing of oral samples (6 experiments, 20.8 Gb). Proteome data consist of 3 SomaScan plasma assays and clinome data of 10 clinical records (9 medical examinations and 1 questionnaire). The 136 sequencing experiments total 9.50 Tb.

LCL, lymphoblastoid cell line; PBMC, peripheral blood mononuclear cell; TSLR, TruSeq synthetic long read.

## Methods

### Ethics statement and sample collection

All sample donors provided written informed consent for participation in this study. Ethical approval was obtained from the Institutional Review Board (IRB) of the Genome Research Foundation (IRB-201307-1 and IRB-201501-1 for KOREF, and IRB-20101202-001 for KPGP). Additionally, the KOREF1 Trio (KOREF1 and his parents) was approved by the Institutional Review Board of the Ulsan National Institute of Science and Technology (UNISTIRB-15-19-A). Blood samples from KOREF1 have been collected longitudinally since 2013, and donors reaffirmed their agreement through a Sample Utilization Consent Form at each collection.

### Genome sequencing

Genomic DNA was extracted from the peripheral blood of a healthy Korean male donor with the Monarch HMW DNA Extraction Kit for Tissue (New England Biolabs, T3060L) and the QIAGEN Blood & Cell Culture DNA Mini Kit (Cat# 13323), following the manufacturers’ protocols. PacBio HiFi sequencing was performed on the Sequel II and Revio platforms. Libraries were prepared with the SMRTbell Express Template Prep Kit 2.0 (Pacific Biosciences, Cat# 100-938-900) with size selection to enrich large-insert fragments. The Sequel II runs yielded 117 Gb of HiFi reads (circular consensus, ccs v6.0.4), and the Revio runs added 152 Gb (ccs v8.0.1), giving 269 Gb in total ∼90×. Ultra-long Oxford Nanopore libraries were prepared with the SQK-ULK114 Ultra-Long DNA Sequencing Kit V14 (Oxford Nanopore Technologies) and sequenced on the PromethION 2 Solo with R10.4.1 flow cells (FLO-PRO114M). Super-accurate basecalling with dorado v0.7.3 produced 922.7 Gb (∼308×). Parental short reads for trio phasing were prepared as paired-end libraries (2×150 bp) with the TruSeq DNA PCR-Free kit (Illumina, Cat# 20015963) and sequenced on the HiSeq X platform, giving ∼40× paternal and ∼37× maternal coverage. Offspring (KOREF1) short reads were generated on the NovaSeq platform to ∼118× coverage.

### Transcriptome sequencing

Transcript evidence for gene annotation was generated with both short-read RNA sequencing and long-read Iso-Seq from whole blood and a B-cell immortalized cell line. For long reads, RNA was processed with the SMARTer PCR cDNA Synthesis Kit (Takara Bio, Cat# 634925) and the SMRTbell Template Prep Kit 1.0 (PacBio, Cat# 100-259-100) and sequenced on the PacBio Sequel platform. Additional whole-blood Iso-Seq libraries were prepared after globin depletion with the GLOBINclear-Human Kit (Ambion/Thermo Fisher, Cat# AM1980) and sequenced on the PacBio Sequel II platform to obtain high-quality full-length transcripts. For short reads, whole-blood libraries were prepared with the TruSeq RNA Library Prep Kit v2 (Illumina, Cat# RS-122-2001/2002) and sequenced on the NovaSeq 6000, and B-cell line RNA was prepared with the TruSeq Stranded mRNA Sample Prep Kit (Illumina, Cat# RS-122-2101/2102) and sequenced on the HiSeq 2500. These short-read datasets provided high-depth coverage for accurate exon-boundary and splice-junction detection.

### Read preprocessing

We prepared high-quality long reads from the raw PacBio HiFi and Oxford Nanopore Technology (ONT) ultra-long (UL) datasets to construct an accurate and gapless genome assembly. For PacBio HiFi, we removed residual adapters with HiFiAdapterFilt^55^ (v2.0.1). We then discarded reads shorter than 1 kb or with a quality value (QV) below 20, retaining ∼87× coverage. For ONT UL reads, we first clipped the low-quality 1.5 kb at each read end. We then kept only reads ≥100 kb with QV ≥15, yielding ∼89× high-quality ultra-long coverage. We processed the parental short reads with fastp^56–58^ (v0.23.4) to derive haplotype-specific k-mers for phasing. We removed adapters, trimmed the first 15 bp and last 5 bp, and discarded reads containing any ambiguous base (N). This left 24× paternal and 21× maternal coverage. We processed the KOREF1 short reads with fastp^56–58^ (v1.0.1) under the same trimming scheme but a relaxed N-base limit of one, reaching 92× coverage.

Transcriptome sequencing data were processed to generate high-quality reads for gene annotation. For PacBio Iso-Seq, raw subreads were converted to circular consensus sequence (CCS) reads with ccs (v6.4.0). Primer sequences were removed with lima (v2.13.0). Full-length non-chimeric (FLNC) reads were then refined and clustered with isoseq (v4.3.0) to obtain high-quality transcript sequences. For Illumina RNA-seq, poly-G tails were removed with fastp^56–58^ (v1.0.1). Adapters were then trimmed, and the first 15 bp and last 5 bp of each read were removed. Reads with more than one ambiguous base (N) were discarded.

### Initial Genome Assembly

Initial assemblies were generated with hifiasm^59–61^ (v0.25.0-r726) in trio mode (--dual-scaf, -- telo-m CCCTAA) from ∼87× PacBio HiFi and ∼89× ONT ultra-long reads (≥100 kb). Parental short-read coverage was 24× paternal and 21× maternal, leaving 0.53 million distinct k-mers at the child’s haploid coverage absent from both parental databases (Figure S1). We therefore replaced hifiasm’s default parental k-mer counting with parental specific k-mers. These k-mers (k = 31) were built from the parental and KOREF1 HiFi reads with meryl (v1.4.1) and Merqury^62^ (v1.3), giving 40.7 million paternal and 28.3 million maternal markers, and converted to yak format (yak^59–61^ v0.1-r69-dirty) as the parental inputs. The assemblies were already near chromosome-scale: 110 paternal and 98 maternal primary contigs spanning 2.92 and 3.04 Gb, contig N50 136.8 and 136.2 Mb, largest contigs 247.5 and 252.7 Mb.

### Genome Scaffolding

Telomeric repeats were identified with tidk^63^ (v0.2.31) using the CCCTAA motif, and a chromosome end was considered telomere-capped when its terminal 5 kb carried more than 100 canonical motifs. Scaffolds missing a telomere at one or both ends were selected for re-scaffolding with parental-binned ONT reads ≥50 kb. After hard-masking the candidate telomeric regions, we re-scaffolded them with ntLink^64,65^ (v1.3.11). RagTag^66,67^ (v2.1.0) was then run in scaffold mode against T2T-CHM13^20^ solely to assign chromosome identity and orientation to the resulting scaffolds. This mode does not modify the input sequence: no base was corrected, broken or inserted from the reference, and every assembled base derives from KOREF1 reads. T2T-CHM13 therefore served only as a coordinate frame for naming and orienting the 23 chromosome-scale scaffolds per haplotype. This completed 23 telomere-to-telomere chromosomes per haplotype, with 44 of 46 ends telomere-capped, except for the paternal chromosome 13 p-arm and the maternal chromosome 15 q-arm.

### Gap Closing

We first restored the missing telomeric ends of paternal chromosome 13p and maternal chromosome 15q by targeted local assembly. HiFi reads were filtered by length and quality (≥1 kb, QV ≥20) and partitioned into paternal, maternal, and unclassified sets with yak^59–61^ triobin (v0.1-r69-dirty). Parental-binned reads were aligned to the opposite-haplotype chromosome ends with Winnowmap2^68,69^ (v2.03) using a 15-mer repetitive database. Because the opposite KOREF1 haplotype is genetically closer than T2T-CHM13^20^, this recovered up to 4.5-fold more reads and reached ∼30× coverage at the target ends. Primary alignments were extracted with samtools^70^ (v1.17), and the reads were locally reassembled with hifiasm^59–61^ (v0.25.0-r726) (--n-hap 1, -l0).

We then closed internal gaps with a modified TGS-GapCloser^71^ (v1.2.1) that improves speed and minimizer-index sampling by aligning reads to assemblies rather than assemblies to reads. Gap closing proceeded in stages with progressively updated inputs. We used haplotype-binned HiFi reads and their re-assemblies with hifiasm^59–61^ (v0.25.0-r726) and Flye^72,73^ (v2.9.6). We retained only assemblies with QV > 50 and Hamming error < 0.05 for gap closing. Remaining gaps were resolved by contig chaining on pairwise contig-to-contig alignments from the earlier assemblies. High-confidence alignments were connected across gaps with a Dijkstra shortest-path search, inserting a 100 bp placeholder gap to prevent overlap errors. These secondary gaps were then closed with trio-binned HiFi reads under stricter filters (identity >97%, alignment ratio >97%). This produced a completely gapless assembly.

### Genome Polishing

Both KOREF1 haplotypes were polished over five iterative rounds, each applying structural-variant (SV) polishing followed by single-nucleotide-variant (SNV) polishing based on the T2T-Polish pipeline^74^. SVs were called from trio-binned ONT and HiFi alignments (VAF >0.4, read support >40%) with Sniffles2^75,76^ (v2.6.3) and refined with Iris^77^ (v1.0.5). Because many heterozygous calls persisted even with haplotype-binned reads, we merged calls across platforms with Jasmine^77^ (v1.1.5) and polished only variants that were homozygous in at least one platform or whose opposite-haplotype locus was reference-homozygous. Where binned-read depth fell below 5, unclassified reads were added and polishing was restricted to those regions to avoid introducing false-positive SVs. Large SVs (>500 bp) were curated manually in IGV^78–81^ (v2.19.6). SNVs were called from HiFi alignments with PEPPER-Margin-DeepVariant^82^ (r0.8) and filtered with bcftools^70^ (v1.22) (biallelic: GQ >30, VAF >0.4; multiallelic: GQ >25, min VAF >0.3, summed VAF >0.7). Heterozygous calls were re-evaluated with trio-binned reads and checked against the opposite haplotype, low-HiFi-depth regions (<5) were supplemented with high-confidence ONT SNVs (VAF >0.8, depth >10), and the set was refined with Merfin^83^ (v1.1) before polishing. Regions lacking both HiFi and ONT alignments remained poorly polished with low QV. We located them by computing QV in 2-kb windows (k=31), longer than the k=21 of the genome-wide quality measurement QV, because a 31-mer recurs less often by chance in a 3 Gb genome and therefore localizes errors inside repeats more specifically, and patched them by anchoring candidate contigs on flanking high-QV sequence, expanding the anchor across three rounds (50, 200, 500 kb) for specificity in repeats. Only contigs with >95% identity to both anchors and a unique placement replaced the low-QV segment. Over the five rounds, the consensus quality value rose from 72.7 to 81.2 (paternal) and from 71.4 to 79.0 (maternal), about one error per 131 Mb and 80 Mb, respectively (Table S1).

### Repeat Annotation

Repetitive elements were annotated and soft-masked on each haplotype with RepeatMasker (v4.2.2) (http://repeatmasker.org) (rmblast, sensitive mode) against the Dfam 3.9 human library^84–87^. Repeats covered 53.17% (paternal) and 53.38% (maternal) of the assembly, dominated by LINEs (20.0% and 20.7%), SINEs (12.9% and 12.8%), LTR elements (8.9%), and satellites (4.9% and 4.8%) (Table S2). Centromeric higher-order repeats (HORs), alpha-satellite, human satellites (HSat1–3), and ribosomal DNA were annotated with the alphaAnnotation pipeline (commit 35e53eb; https://github.com/kmiga/alphaAnnotation), resolving 266.3 (paternal) and 252.9 Mb (maternal) of centromeric satellite, including 64.0 and 67.5 Mb of active α-satellite HOR across the 23 centromeres of each haplotype (Table S3). Segmental duplications were identified on the soft-masked genome with BISER^88^ (v1.4) after removing self-alignment pairs with ≥50% mate-region overlap. Recent SDs (>90% identity) spanned 97.5 Mb (3.35%, paternal) and 103.8 Mb (3.43%, maternal). Extending to 70% identity added ancient SDs, reaching 216.8 Mb (7.44%) and 218.8 Mb (7.22%) (Table S3).

### Gene Annotation

Gene models were annotated with the Comparative Annotation Toolkit (CAT v2.2.1; Augustus CGP and PB modes) over GRCh38-to-haplotype whole-genome alignments from Cactus-Pangenome (v9.1.2) ^89^, using GRCh38 GENCODE v49 as the reference^16^. Transcript evidence comprised 675,861 full-length non-chimeric Iso-Seq reads (11 SMRT cells, blood and cell line; spliced-aligned with minimap2^90,91^ v2.29) and 80.6 million RNA-seq pairs (blood 45.2 M, cell line 35.3 M; STAR^92^ v2.7.10b two-pass, 63.1% and 87.9% uniquely mapped). Transcripts that CAT flagged as unknown-likely-coding were triaged for coding evidence: a Swiss-Prot hit^93^ (DIAMOND^94,95^ v2.1.22, E ≤ 1×10), an InterProScan^96,97^ (v5.77-108.0) domain (AntiFam-spurious removed), or Salmon-quantified expression^98^. Those without protein evidence that were lowly expressed (TPM < 0.5) and short (CDS < 100 bp) were removed, leaving 1,552 (651 paternal, 901 maternal). We resolved all 1,552 by mapping their gene regions to T2T-CHM13^20^ and GRCh38^16^ (minimap2 asm20) and transferring the overlapping reference gene, recovering 675 as protein-coding; each was then counted by its curated biotype rather than as unknown-likely-coding. Genes absent from CAT were supplemented with Liftoff^99^ (v1.6.3) from CHM13^20^ CAT/Liftoff (GENCODE v35) and GRCh38^16^ (GENCODE v49). The final annotation held 78,316 (paternal) and 81,174 (maternal) gene features, including 19,431 and 20,404 protein-coding genes, 34,708 and 35,412 lncRNA genes, and 15,056 and 15,604 pseudogenes (Table S3).

### Candidate cis-regulatory element (cCRE) annotation

We projected the ENCODE SCREEN V4 human cCRE registry^100^ (2,348,854 elements across eight classes: PLS, pELS, dELS, CA-CTCF, CA-H3K4me3, CA-TF, CA, and TF) from GRCh38 onto both KOREF1 haplotypes. Projection used a chain built from GRCh38 to each KOREF1 haplotype from per-chromosome minimap2^90,91^ (v2.29) asm5 alignments with the cs tag. paf2chain (v0.1.1) converted each alignment to a chain, and the UCSC kentUtils v483 suite rescored and netted it: chainScore rescored the chain against the two-bit sequences with the default matrix because paf2chain emits a constant score that collapses the net hierarchy, then chainSort, chainPreNet, chainNet, and netChainSubset produced the final chain. CrossMap^101^ (v0.7.0) lifted the cCRE intervals. CrossMap fragments an element wherever it spans a chain-block boundary, so we merged the fragments of each SCREEN accession back into one interval. This affected 106,629 elements in the paternal and 108,306 in the maternal haplotype (about 4.7 and 4.6 %); the remaining 95 % lifted as a single block. The maternal haplotype retained 2,343,133 cCREs (99.8 %) and the paternal haplotype 2,280,991 (97.1 %). The paternal deficit is the 65,915 GRCh38 chrX elements with no target in the chrY-bearing haplotype, and the reciprocal 3,401 chrY elements account for most of the maternal loss. Excluding the haplotype-absent sex chromosome, unmapped elements fall to 0.08 to 0.10 % per haplotype.

### CpG island (CGI) annotation

We annotated 48,394 CpG islands on the paternal haplotype (36.2 Mb, median 567 bp) and 49,977 on the maternal (37.7 Mb, median 573 bp), each at mean 58.3 % GC and mean observed-to-expected CpG 0.74. Islands were called *de novo* by the Gardiner-Garden and Frommer criteria as a Takai and Jones sliding-window scan^102^: a 200 bp window advanced one base at a time, and a merged window run was kept when it spanned at least 200 bp, held at least 10 CpG dinucleotides, and reached at least 50 percent GC and an observed-to-expected CpG ratio of at least 0.6 over its full length. We scanned the soft-masked assemblies and counted only uppercase C and G, which removed the 2.2-fold over-calling of the unmasked scan (115,373 paternal islands) from constitutively methylated repeats; overlapping islands were merged with statistics recomputed from sequence. We then annotated the 2 kb flanking each island as its shore, giving 79,741 shores on the paternal haplotype and 82,149 on the maternal.

### Chromosome Y annotation

Chromosome Y was annotated into pseudoautosomal, X-transposed, X-degenerate, ampliconic and heterochromatic regions using T2T-CHM13 (v2.0) chrY (same as HG002 chrY) (62,460,029 bp) as the reference^103,104^. Eleven anchor elements (PAR1, PAR2, two X-degenerate and two X-transposed segments, SAT1, SAT2, CEN, DYZ17, DYZ19) were aligned to the KOREF1 chromosome one at a time with minimap2^90,91^ (v2.29) (-cx lr:hqae --cs --secondary=yes -N 20, 30– 200 kb minimum alignment). Intervals between anchors took their flanking label. DYZ19 was then located by a search restricted to the ampliconic zone, and the X-degenerate segments of the male-specific region were transferred by whole-chromosome alignment. The boundaries of the seven named palindromes AMPL1–AMPL7 were set from the positions of their flanking marker sequences (minimap2^90,91^ -cx asm5, MAPQ ≥ 20). Yq12 heterochromatin was delineated by nhmmer^105–107^ (HMMER v3.4) against DYZ1 and DYZ2 satellite consensus, and its proximal boundary was placed at the start of the terminal contiguous DYZ array, scored in 50 kb windows. The annotation covers the chromosome completely, 47,256,451 bp in 37 intervals across 17 region types, of which 19.62 Mb (41.5%) is Yq12 heterochromatin, 10.48 Mb (22.2%) ampliconic, 8.79 Mb X-degenerate, 3.35 Mb X-transposed and 2.21 Mb pseudoautosomal. The AZFc b2/b3 inversion was genotyped from the orientation of 14 CHM13 probes (minimap2 -cx asm5 --secondary=yes -N 5), read against the P3 spacer, which lies outside the inversion and maps uniquely at MAPQ 60. KOREF1 carries the inversion, at 97% confidence across four diagnostic amplicon arms. KOREF1 has the shortest chromosome Y of the four telomere-to-telomere chromosomes compared, 47.3 Mb against 51.5 Mb for T2T-YAO, 65.7 Mb for CN1 and 62.5 Mb for CHM13. The euchromatic male-specific region is 25.43 Mb in KOREF1 and 24.8 to 25.4 Mb in the other three, in the same class order, so the 18.4 Mb range in total length lies entirely in Yq12 heterochromatin, 19.6 to 37.6 Mb (Figure S2).

### Genome Quality Evaluation

Terminal telomeric arrays was quantified with tidk^63^ (v0.2.31), counting canonical CCCTAA repeats at every chromosome end. Genome Continuity Inspector (GCI) ^108^ (v1.0) scored per-haplotype continuity. It used HiFi (≥1 kb, QV ≥20) and ONT (≥10 kb, QV ≥15) reads aligned to each haplotype with both Winnowmap2^68,69^ (v2.03) and minimap2^90,91^ (v2.29). Passing the two alignments together, GCI kept only reads placed consistently by both aligners.

HMM-Flagger^35,109^ (v1.1.0) assessed structural reliability. HiFi and ONT reads were aligned to the diploid assembly (both haplotypes, 46 chromosomes) with Winnowmap2^68,69^ (v2.03). SecPhase (v0.4.4) then reassigned multi-mapping reads to their correct haplotype. Flagger^35,109^ classified every base as reliably assembled or as an error component (erroneous, duplicated, or collapsed). Centromeres, sex chromosomes, and segmental duplications were supplied as bias-prone priors. ModDotPlot^110^ (v0.9.8), in comparison mode, computed the pairwise sequence identity between each KOREF1 HOR array and its corresponding T2T-CHM13 array. Bionano optical maps (Bionano Solve v3.8.3.1) validated the ribosomal DNA arrays, which collapse in read-only assemblies. Hi-C contact maps (HiC-Pro^111^ v3.1.0, HiCExplorer^111–114^ v3.6) confirmed chromosome-scale continuity from their single-diagonal pattern.

We scored single-copy gene completeness with compleasm^115^ (v0.2.7) against the primates lineage (OrthoDB odb12^116,117^). NucFlag^74,118,119^ (v0.3.7) examined the secphase-corrected diploid long-read pileups. It flagged misjoins, collapses, and false duplications, keeping only structural misassemblies. VerityMap^120^ (v2.0.0) verified satellite copy number by anchoring rare k-mers within the centromeric-satellite (cenSat) arrays. CRAQ^121^ (v1.10) separated genuine structural errors from the soft-clips left by heterozygous variants. It computed the genome-wide Assembly Quality Index (AQI) and mapped clip-based regional (CRE) and structural (CSE) errors along each chromosome. We then intersected these error regions with the gene and cis-regulatory element (cCRE) annotations to obtain per-class error rates. Merqury^62^ (v1.3) computed consensus quality value (QV) and k-mer completeness from KOREF1 hap-mers. Per-chromosome scores for every metric are reported for both haplotypes in Table S4.

### Mitochondrial Genome Assembly

The mitochondrial genome was assembled directly from the PacBio HiFi sequencing data (51 x coverage, two SMRT flow cell) using the MitoHiFi pipeline^122–125^ (v3.2.1) with default parameters. We used the revised cambridge reference sequence (rCRS) of the human mitochondrial DNA as reference^126^. The resulting KOREF1 mitochondrial genome was assembled as a single circular contig of 16,570bp, representing the complete mitochondrial sequence with no gaps. All 37 mitochondrial genes, including protein-coding genes, tRNAs and rRNAs were also annotated.

## Data Records

The KOREF1 assemblies, annotations, and multi-omic reference dataset, comprising raw sequencing reads, processed data, and analysis outputs, are accessible through a data portal with an integrated genome browser at https://koreanreference.org. All sequencing data generated for the KOREF1 individual have been deposited in NCBI under BioProject PRJNA1037546, and the diploid assemblies under BioProject accessions PRJNA1508392 and PRJNA1508401. The paternal and maternal genome assemblies of KOREF1-G-TTAGGA have additionally been deposited in the Korea BioData Station (K-BDS) under Project ID KAP242509.

## Technical Validation

### Telomere-to-Telomere Accurate and Gapless Genome Assembly

We present KOREF1-G-TTAGGA, a Telomere-to-Telomere Accurate and Gapless Genome Assembly (TTAGGA) of KOREF1. The diploid assembly (44 + XY + mtDNA) comprises a 2.91 Gb paternal (NG50 = 146.35Mb) and a 3.03 Gb maternal haplotype (NG50 = 155.66Mb) (Figure 2A and 2B, Table S1). Every chromosome was reconstructed as a single gapless contig, with telomeric repeat arrays (1,049 ± 459 repeats, 6.3 ± 2.8 kb) capping every terminus (Figure S3). That completeness closes the 997 gaps, totaling 338 Mb, that remained in the previous KOREF1 release^45^ (Figure S4). Genome Continuity Inspector (GCI) ^108^ scored 84.92 and 80.78 for the paternal and maternal haplotype, respectively. These exceeded those of Han Chinese reference genome CN1 (77.90 and 66.79) ^29^ and approached T2T-CHM13 (87.04) ^20^, the first complete human reference genome (Figure S5). A complete genome, however, is not just unbroken but correct in both structure and sequence.

**Figure 2.**
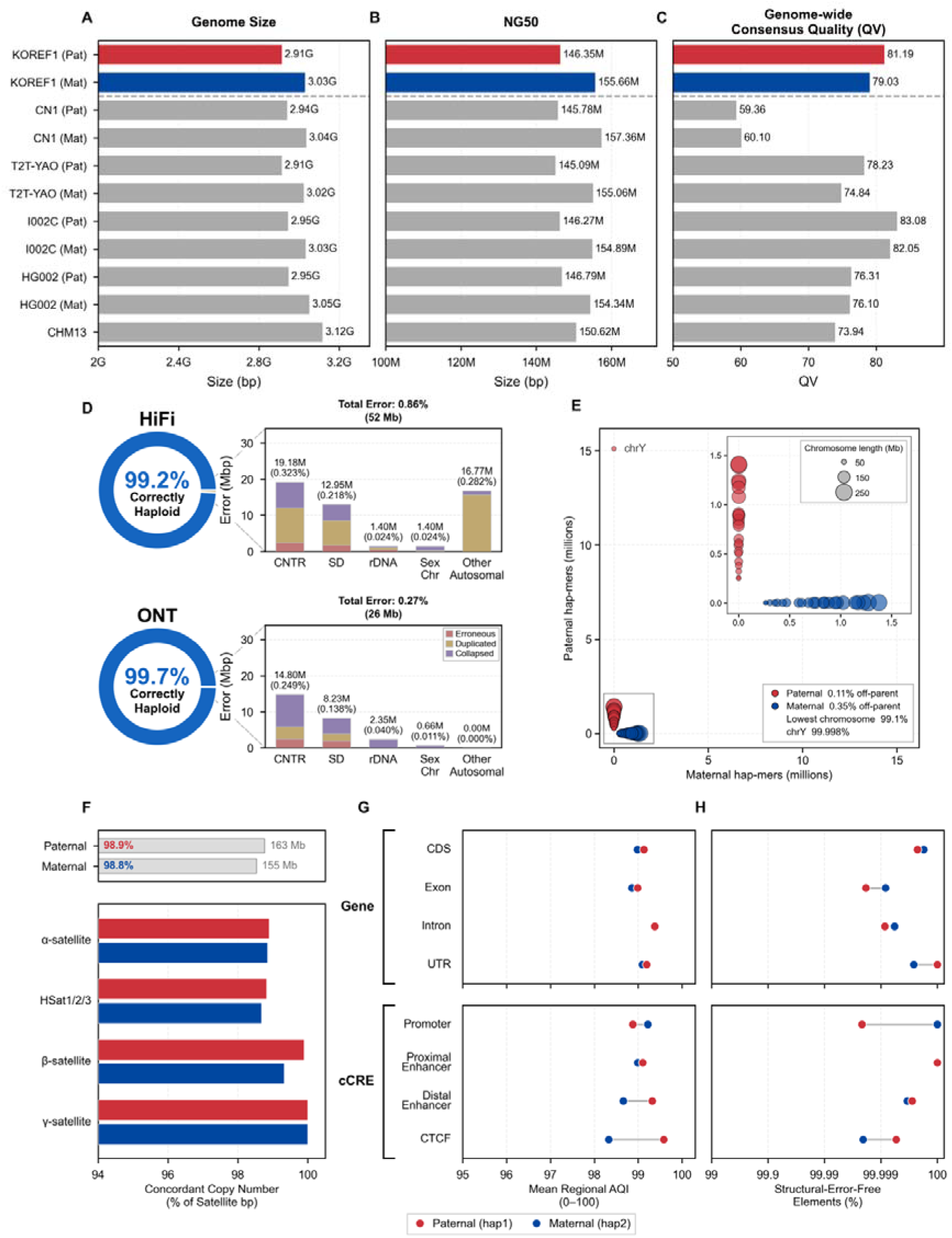
Continuity, structural integrity, and base accuracy of the KOREF1-G-TTAGGA assembly. (A–C) Assembly statistics for the KOREF1 paternal (red) and maternal (blue) haplotypes, compared with published human telomere-to-telomere (T2T) assemblies (grey): T2T-CHM13^20^, HG002^103,104^, CN1^29^, T2T-YAO^128,129^, and I002C^130^. Bars show (A) total assembled size, (B) contig NG50, and (C) genome-wide consensus quality (Merqury^62^ QV, Phred-scaled; higher is better). QV values for the comparator assemblies are taken from their respective publications. (D) Per-base reliability estimated with Flagger^35,109^ from HiFi (top) and ONT (bottom) long-read alignments (winnowmap2^68,69^), across the diploid assembly. Each donut gives the fraction of the assembly classified as correctly assembled ("Correctly Haploid"); the remainder is partitioned into Erroneous, Duplicated, and Collapsed bases. Adjacent stacked bars show the error (Mbp, and % of the genome in parentheses) contributed by each region class: CNTR (centromere), SD (segmental duplication), rDNA (ribosomal DNA), Sex Chr (sex chromosomes), and Other Autosomal. Bar titles give the total flagged error per platform. (E) Trio hap-mer content of each chromosome, plotted as maternal against paternal parent-specific k-mers called by Merqury ^62^ from the parental short reads, with marker area proportional to chromosome length. A correctly phased chromosome carries hap-mers from one parent only and therefore lies on an axis, whereas a chimeric one would fall towards the diagonal. Axes span the full range so that chromosome Y is on scale (15,104,277 paternal against 318 maternal hap-mers); the grey box marks the autosomes and chromosome X, redrawn in the inset on the same marker-size scale. The key gives the fraction of assigned hap-mers derived from the opposite parent, the lowest per-chromosome purity, and chromosome Y. (F) Satellite copy-number concordance (VerityMap^120^). Top bars give the genome-wide fraction of paternal (163 Mb) and maternal (155 Mb) satellite sequence with concordant copy number; the lower bars resolve this by satellite family (α-satellite, HSat1/2/3, β-satellite, γ-satellite). (G, H) Functional-element accuracy from CRAQ^121^ across gene features (CDS, exon, intron, UTR) and candidate cis-regulatory elements (cCRE: promoter, proximal enhancer, distal enhancer, CTCF), for the paternal (hap1, red) and maternal (hap2, blue) haplotypes: (G) Mean regional AQI: the CRAQ window score (∼3 Mb windows) averaged over every base of each annotation class, and (H) Percentage of elements in each class with no overlapping CRAQ error interval. (99–100%). TTAGGA, Telomere-to-Telomere Accurate and Gapless Genome Assembly; NG50, contiguity metric; QV, consensus quality value; hap-mer, parent-specific k-mer; AQI, Assembly Quality Index; cCRE, candidate regulatory element.

Even where mis-phasing and fragmentation typically arise, KOREF1-G-TTAGGA remains structurally intact. Genome-wide, read-depth reliability estimation (Flagger) ^35,109^ classified at least 99.15% of the assembly as reliably assembled and correctly haploid, with minimal loss of fidelity in structurally complex regions, exceeding 95% within centromeres and 97% within segmental duplications (Figure 2D and Figure S6, Table S5). Trio hap-mers assigned every chromosome to a single parental haplotype, with off-parent content of paternal 0.11% and maternal 0.35%, no chromosome below 99.1% purity, and the paternal chromosome Y at 99.998% (Figure 2E). All 46 α-satellite higher-order repeat (HOR) arrays were assembled at full length (145.8 Mb, ≥96% Flagger read-supported), and highly homologous to T2T-CHM13^20^ (Figure S7 and Table S3). The collapse-prone rDNA arrays were spanned end-to-end by Bionano optical maps, structurally validating their full-length assembly (Figure S8). Hi-C contact maps resolved each chromosome as a single continuous diagonal with no discrete off-diagonal blocks, confirming chromosome-scale integrity of both haplotypes (Figure S9 and S10).

Both haplotypes showed completeness and correctness at the locus level. Their gene space was complete, carrying the full set of primate single-copy orthologs (BUSCO) ^115–117^ with no fragmented genes, on par with the T2T-CHM13 reference^20^ (Figure S11). NucFlag^74,118,119^, examining the long-read pileups against each haplotype, scored 99.92% and 99.91% concordance (Figure S12 and S13). Rare k-mer anchoring (VerityMap) ^120^ confirmed accurate satellite copy number across 98.9% and 98.8% of the paternal and maternal satellite sequence, even in the most repetitive arrays (Figure 2F and S14). Clipping Reveals Assembly Quality (CRAQ) ^121^, which separates structural errors from the clips that heterozygous variants leave, set the Assembly Quality Index (AQI) at 99.77 and 99.69, far surpassing the reference-quality threshold of 90. The mean regional AQI ranged from 98.3 to 99.6 across gene and cCRE classes, and more than 99.99% of gene and cCRE sequence was free of structural error (Figure 2G and 2H, Figure S15 and S16).

KOREF1-G-TTAGGA further achieved high accuracy at the base level. The assembly recovered nearly all sequence content present in the reads, with diploid k-mer completeness of 99.85% and parent-specific k-mer (hap-mer) completeness of 99.78% (paternal) and 99.85% (maternal) (Table S1). Structural error called concordantly by at least two of Flagger^35,109^, GCI^108^, CRAQ^121^ and NucFlag^74,118,119^ covered 13.5 Mb across 153 loci (0.23% of the assembly, 77 paternal and 76 maternal), of which 12.8 Mb lay in highly repetitive, near-identical sequence, mostly satellite arrays and segmental duplications (Figure 3A and 3B). Merqury-based consensus quality (QV) scores^62^ were 81.19 (paternal) and 79.03 (maternal), representing fewer than one error per 131Mb and 80Mb (Table S1). These exceeded T2T-CHM13 (73.9) ^20^, HG002 (paternal: 76.10, maternal: 76.31) ^103,104^ and the East Asian telomere-to-telomere assemblies, including CN1 (paternal: 59.36, maternal: 60.10) ^29^ and T2T-YAO (paternal: 78.23, maternal: 74.84) ^128,129^ (Figure 2C). Per-chromosome QV averaged 82.0 ± 6.3 (paternal) and 80.2 ± 8.2 (maternal) among chromosomes and only 0.0025% of 10kb windows (15 loci, 122 kb) fell below QV 40 (Figure 3A and 3B). KOREF1-G-TTAGGA thus achieves one of the highest base-level accuracies among the East Asian T2T assemblies.

**Figure 3.**
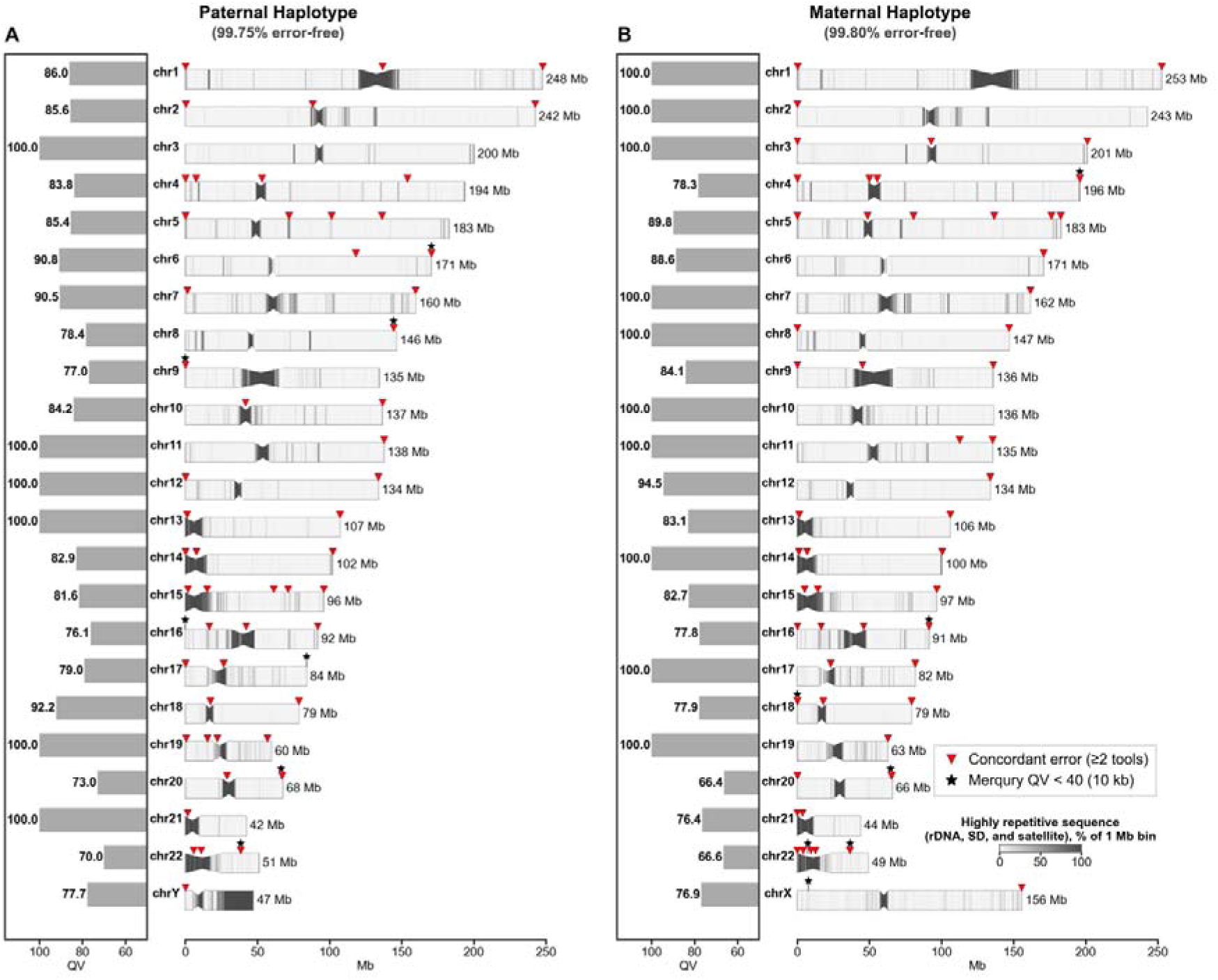
Chromosome-scale integrity and consensus quality of the KOREF1-G-TTAGGA diploid assembly. Whole-genome ideograms of the (A) paternal (hap1; autosomes and chrY) and (B) maternal (hap2; autosomes and chrX) haplotypes. For each chromosome, the bar at left gives the per-chromosome consensus quality (Merqury^62^ QV, k = 21; axis reversed, higher quality toward the left, bars truncated at QV 50). Sixteen chromosomes carry no assembly-only k-mer and are plotted at 100. The ideogram at right shows the chromosome drawn to a common 250 Mb scale with its length labelled at the right end. Chromosome bodies are shaded by the fraction of each 1 Mb bin that is highly repetitive, near-identical sequence, taken as rDNA, satellite (AlphaAnnotation; https://github.com/kmiga/alphaAnnotation, transition regions included) and segmental duplication (BISER^88^) and made mutually exclusive in that order. The constriction marks the active α-satellite higher-order repeat array. Red triangles above each chromosome denote assembly-error loci, defined as 100 kb bins independently called by at least two quality-control methods (Flagger^35,109^ erroneous and collapsed blocks, Genome Continuity Inspector^108^, CRAQ^121^, and NucFlag^74,118,119^ mis-join, collapse and false-duplication calls); adjacent bins are merged into 153 loci, drawn as 107 triangles after merging loci within 3 Mb that would overlap at this scale, and requiring two-method concordance removes tool-specific artifacts (Methods). Black stars mark the 15 non-overlapping 10 kb Merqury^62^ windows falling below QV 40. Panel subtitles give the fraction of each haplotype that is structurally error-free under this criterion (paternal 99.75%, maternal 99.80%). QV, consensus quality value; hap1/hap2, paternal/maternal haplotype; SD, segmental duplication; HOR, higher-order repeat.

### Where Accuracy Ends: Reads That Validate Themselves and Overestimated Novelty

KOREF1-G-TTAGGA is gapless and accurate across the genome, but a small amount of error remains unresolved within near-identical repeat arrays. Structural error called concordantly by at least two of Flagger^35,109^, GCI^108^, CRAQ^121^ and NucFlag^74,118,119^ amounts to 13.48 Mb, 0.227% of the 5.94 Gb diploid assembly (Figure 3). Nearly three-quarters of it sits on the five acrocentric chromosomes (9.77 Mb, 72.5%), and chromosome 22 alone accounts for 4.42 Mb (32.7%) (Figure 3 and Table S6). Little of this sequence carries functional elements. Most intervals contain no gene at all (1,891 of 2,510, 75.3%), and 99.975% of coding regions and 99.989% of candidate cis-regulatory elements (cCREs) lie outside any error interval. The 619 intervals that do overlap genes are dominated by the rDNA transcription units of the acrocentric short arms and the microRNAs embedded in the rDNA repeat, which account for nearly half of their sequence. The rest falls in paralogue clusters, including PKD1 with its six chromosome 16 pseudogenes and the POTE and NPIP families, and the CT47 macrosatellite on Xq24.

The figures above do not escape self-validation. The four quality assessments realign the HiFi and ONT reads used to build the assembly. Merqury^62^ QV likewise measures how far the k-mers of the assembly are contained in the k-mer set of those reads. Agreement between them therefore cannot exceed what those reads contain. Accurate assessment requires an independent sequencing dataset or a different technology such as optical mapping or Hi-C. Those technologies, however, are weakest exactly where the errors concentrate: highly repetitive sequence holds 95.3% of the called error bases (Figure 3). Our optical maps resolve the rDNA arrays (Figure S8) and cover 96.4% of unique sequence, but the active alpha-satellite arrays carry so few restriction enzyme sites that only 2.3% is covered, and Hi-C reads place almost no contacts there. Inside the arrays, even long-read alignment depends on rare k-mers as anchors, and VerityMap^120^ can flag errors only where such anchors exist (Figure S9). As a result, 52 of the 153 concordant error regions (Figure 3) have no support beyond the assembly’s own reads (8.7 of their 25.6 Mb). Nearly all telomere-to-telomere assembly validated with its own reads carry this gap, and this one is no exception.

Gene annotation is least accurate at the same loci. Liftover from a reference annotation finds low accurate alignment in segmental duplications and near centromeres and telomeres, and *ab initio* prediction fills the gap with gene models that duplicate genes annotated elsewhere in the genome. Of 1,552 loci initially labelled unknown likely coding in this assembly, 1,488 (95.9%) align to CHM13 or GRCh38 once the genomic locus, rather than the predicted protein, is mapped back, and 1,332 (85.8%) land on a gene those references already annotate. Protein homology alone would have missed nearly half of them, since 312 of the 675 recovered protein-coding loci had no homology hit. The 64 loci with no reference alignment are dominated by repeat-derived models: 29 rest on LINE-1, 6 on satellite or simple repeats, and 6 more are ORFs that AntiFam flags as spurious. None is a validated novel protein-coding gene. These regions also differ between individuals. rDNA arrays vary severalfold in copy number, alpha-satellite arrays differ by megabases in length, and the POTE and NPIP families gain and lose copies between genomes. The unknown likely coding loci themselves come from such families: REXO1L, a copy-number-variable pseudogene array on chromosome 8, accounts for 189 of the 1,552, and NBPF and TSPY recur. At these loci, assembly artifact is confounded with genuine biological variation, and a sequence absent from the references can arise from either. Novel genes, novel variants and apparent diversity reported there are therefore easily overestimated, and each candidate needs verification such as mapping the locus back to a reference genome and confirming that no orthologue exists.

## Usage Notes

Bioinformatic data formats have converged on community standards, yet public repositories have been slower to accommodate them. Raw sequencing reads are readily archived, but the processed and analyzed data that constitute the substance of a study have no comparable home.

We therefore built the KOREF Data Portal (https://koreanreference.org) to release every layer of the resource ourselves. Following the open model of the Personal Genome Project (PGP), the portal serves the genome, the multi-omic data, and all intermediate and analyzed outputs without authentication, under explicit versioning so that each release remains citable and reproducible.

To make the resource usable by researchers who do not work at the command line and programmatic access, the portal embeds a genome browser through which the KOREF1-G-TTAGGA sequence and its functional annotations can be inspected directly. The portal was implemented in Next.js, React, and TypeScript, with a PostgreSQL backend and a Tailwind CSS front end. The genome browser was built on JBrowse 2 React components^127^, enabling navigation between the paternal and maternal assemblies and display of genome annotations, structural features, and quality-assessment tracks. Openly releasing not only the genome but every layer of its interpretation is, we believe, how precision medicine moves from principle to practice.

## Data Availability

The data underlying this article are openly available without restriction. Sequencing data generated from the KOREF1 individual are archived at NCBI under BioProject PRJNA1037546, and the diploid assemblies under accessions PRJNA1508392 and PRJNA1508401. The paternal and maternal haplotypes of KOREF1-G-TTAGGA are further archived at the Korea BioData Station (K-BDS) under Project ID KAP242509. A centralized portal at https://koreanreference.org hosts an integrated genome browser and provides bulk access to every component of the resource, including assemblies, annotations, raw reads, processed files, and analysis outputs.

## Code Availability

Analyses were performed using publicly available software as described in the Methods, with versions and parameters specified therein. Custom scripts and analysis workflows are available at https://github.com/korean-genomics-center/KOREF1-G-TTAGGA.git and archived at Zenodo (DOI: 10.5281/zenodo.22064223).

## Author Contributions

Jo.B. conceived and initiated the Korean Standard Reference Genome Project. Ji.B. and Y.K. designed the assembly project. Y.K. assembled the paternal and maternal KOREF1-G-TTAGGA, and Y.C. assembled the mitochondrial genome. S.P. and C.Y. performed molecular library preparation and sequencing. B.D. and J.L. constructed the KOREF1 iPSC line. Ji.B., D.-H.S., K.A., and H.C. organized and curated the KOREF1 multi-omic dataset. Ji.B. and Y.K. jointly performed quality control, evaluation, annotation, and visualization of the assemblies. Ji.B. developed the data portal and genome browser. Ji.B. wrote the original draft. H.L. managed IRB approval and budget administration. H.J. and Jo.B. supervised the project and secured funding. Y.K., D.-H.S., K.A., H.C., D.P., S.J., Y.J., J.J., S.J, J.O.Y., J.-H.K., Y.S.C., J.K., K.S.C., C.G.K., B.D., J.L., S.L., H.J., Jo.B. reviewed and edited the manuscript. All authors read and approved the final manuscript.

## Acknowledgement

We thank George M. Church for his contribution to the initiation of the Korean Genome Project. We also thank all participants of the Korean Genome Project and the Korean Personal Genome Project (KPGP), BioBigData.Korea, and all members of the Korean Genomics Center (KOGIC) and Korean Bioinformation Center (KOBIC). We are especially grateful to the former members of KOGIC, whose work over the years laid the groundwork for the Korean Genome Project. We also acknowledge the Korea Institute of Science and Technology Information (KISTI) for providing the Korea Research Environment Open NETwork (KREONET). The authors used Claude (Anthropic) and ChatGPT (OpenAI) for language editing and to assist with code development for data analysis; All generated text and code were reviewed, tested, and validated by the authors, who take full responsibility for the content of this work.

