## Supplementary Figures S1-S16 for "Telomere-to-Telomere Accurate and Gapless Korean Standard Reference Genome"

**Supplementary Information**

**Index of Supplementary Figures**

Supplementary Figure S1. HiFi read k-mer spectrum and trio hap-mer inheritance.	2

Supplementary Figure S2. ChrY architecture of four telomere-to-telomere assemblies.	3

Supplementary Figure S3. Telomeric repeat arrays across KOREF1-G-TTAGGA chromosomes.	4

Supplementary Figure S4. Comparison of KOREF1-G-TTAGGA with the previous KOREF assembly (KOREF-S1 v2.1).	5

Supplementary Figure S5. Genome continuity assessed by Genome Continuity Inspector (GCI).	6

Supplementary Figure S6. Haplotype-resolved assembly reliability from HMM-Flagger.	7

Supplementary Figure S7. Centromere and satellite organization.	8

Supplementary Figure S8. Optical-map validation of ribosomal DNA (rDNA) arrays.	9

Supplementary Figure S9. Genome-wide Hi-C contact maps.	10

Supplementary Figure S10. Chromosome-wise Hi-C contact maps of the paternal and maternal KOREF1-G-TTAGGA.	11

Supplementary Figure S11. Gene completeness assessment of KOREF1-G-TTAGGA and other high-quality human genome assemblies.	12

Supplementary Figure S12. Read-pileup assembly evaluation (NucFlag) – paternal haplotype.	13

Supplementary Figure S13. Read-pileup assembly evaluation (NucFlag) – maternal haplotype.	14

Supplementary Figure S14. Satellite-array validation by rare k-mer mapping (VerityMap).	15

Supplementary Figure S15. Assembly quality by functional element class (CRAQ).	16

Supplementary Figure S16. Per-chromosome assembly quality (CRAQ).	17

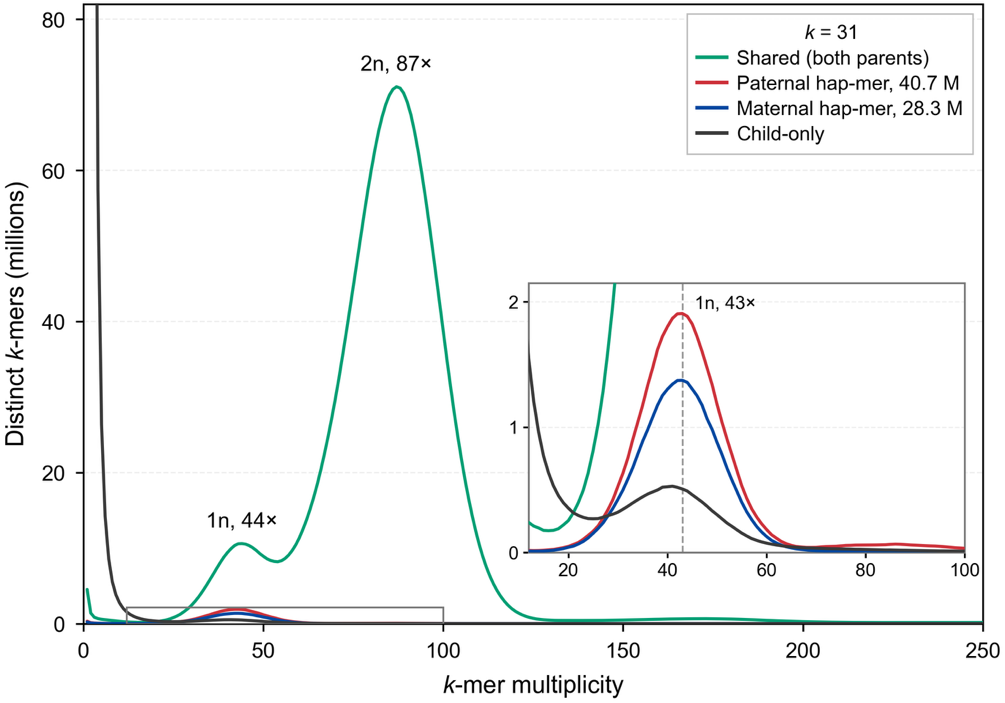

**Supplementary Figure S1. HiFi read k-mer spectrum and trio hap-mer inheritance.** Every 31-mer of the KOREF1 HiFi read set was assigned against meryl databases built from Illumina reads of both parents. Counts are distinct k-mers. K-mers present in both parents resolve a heterozygous peak at 44× and a homozygous peak at 87×, placing HiFi k-mer coverage at 87×, and no further mode is present. K-mers absent from both parents are confined to the low-multiplicity spike at the left edge. The inset magnifies the boxed strip. Paternal and maternal hap-mers both peak at 43×, the haploid coverage, so each parental haplotype is represented at full depth. The child-only curve reaches 0.53 million there, 28% of the paternal peak. Legend totals are distinct hap-mers at or above the meryl inheritance cut-off, 11 for paternal and 13 for maternal.

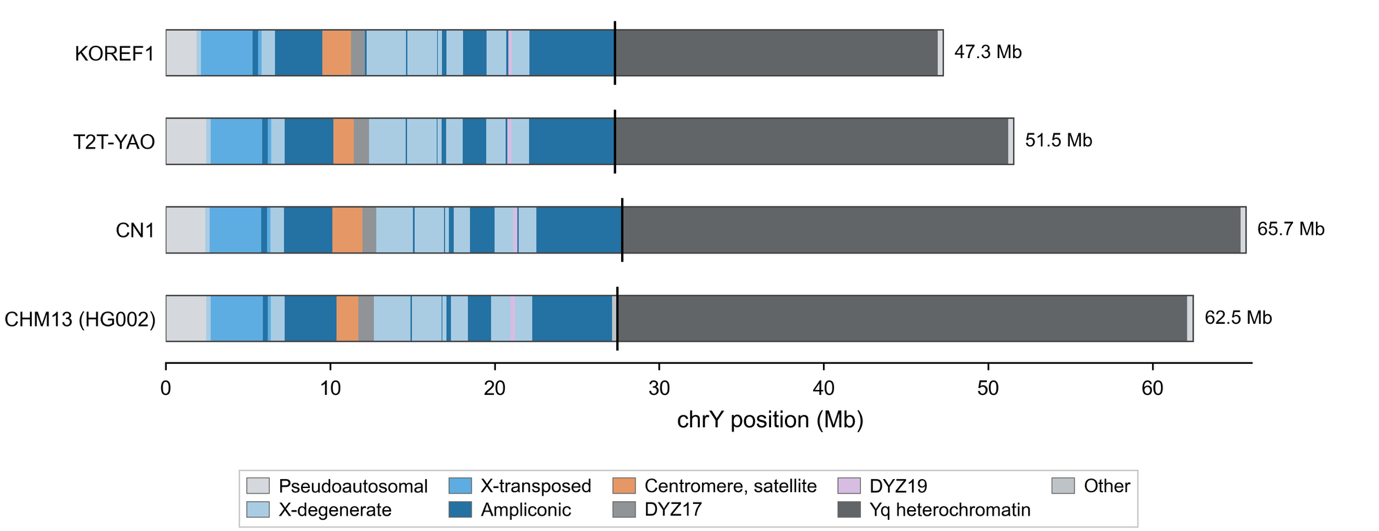

**Supplementary Figure S2. ChrY architecture of four telomere-to-telomere assemblies.** Rows, top to bottom: KOREF1 (hap1, paternal), T2T-YAO, CN1, and CHM13, whose chrY was contributed by HG002, drawn on a common scale. Classes for KOREF1, T2T-YAO and CN1 were transferred from the CHM13 chrY sequence-class annotation by per-element minimap2 alignment, with the named palindromes AMPL1 to AMPL7 placed by nhmmer and pooled here into the ampliconic class. CHM13 carries its own annotation. The vertical tick on each bar marks the start of the terminal DYZ1/DYZ2 array, called from the sample's own DYZ coverage. Total chrY length spans 47.3 to 65.7 Mb, and the euchromatic male-specific region accounts for 24.8 to 25.4 Mb of that in every assembly, in the same class order. The remaining length is Yq heterochromatin, 19.6 to 37.6 Mb, a 1.9-fold range, and its boundary falls within 0.45 Mb of the same position in all four.

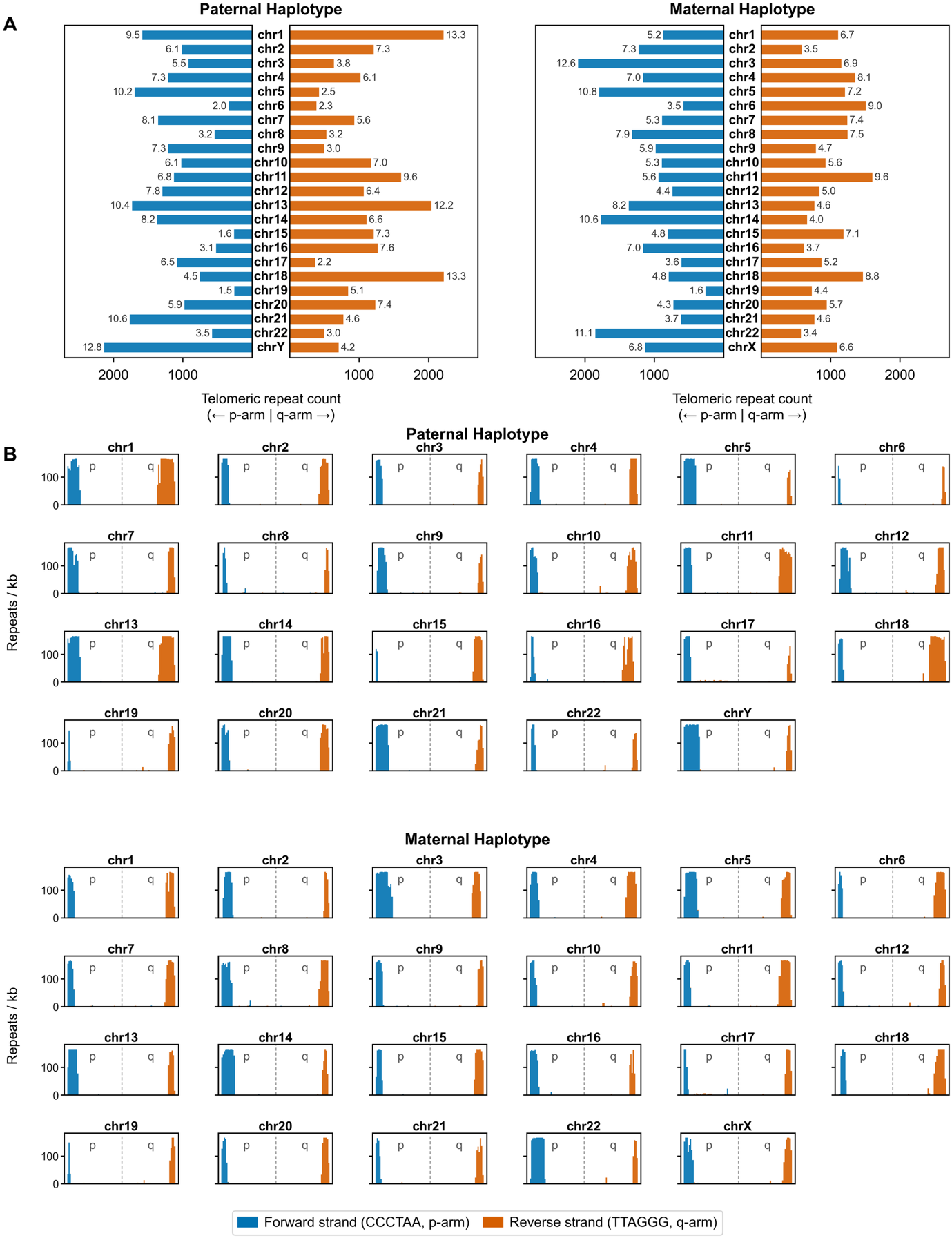

**Supplementary Figure S3. Telomeric repeat arrays across KOREF1-G-TTAGGA chromosomes.** (A) Number of canonical telomeric repeats at the p-arm (left) and q-arm (right) terminus of each chromosome for the paternal and maternal haplotypes; the value beside each bar denotes the telomeric array length (kb). (B) Telomeric repeat density (repeats/kb) along the terminal regions of every chromosome, showing forward-strand (CCCTAA; p-arm, blue) and reverse-strand (TTAGGG; q-arm, orange) motifs for the paternal and maternal haplotypes.

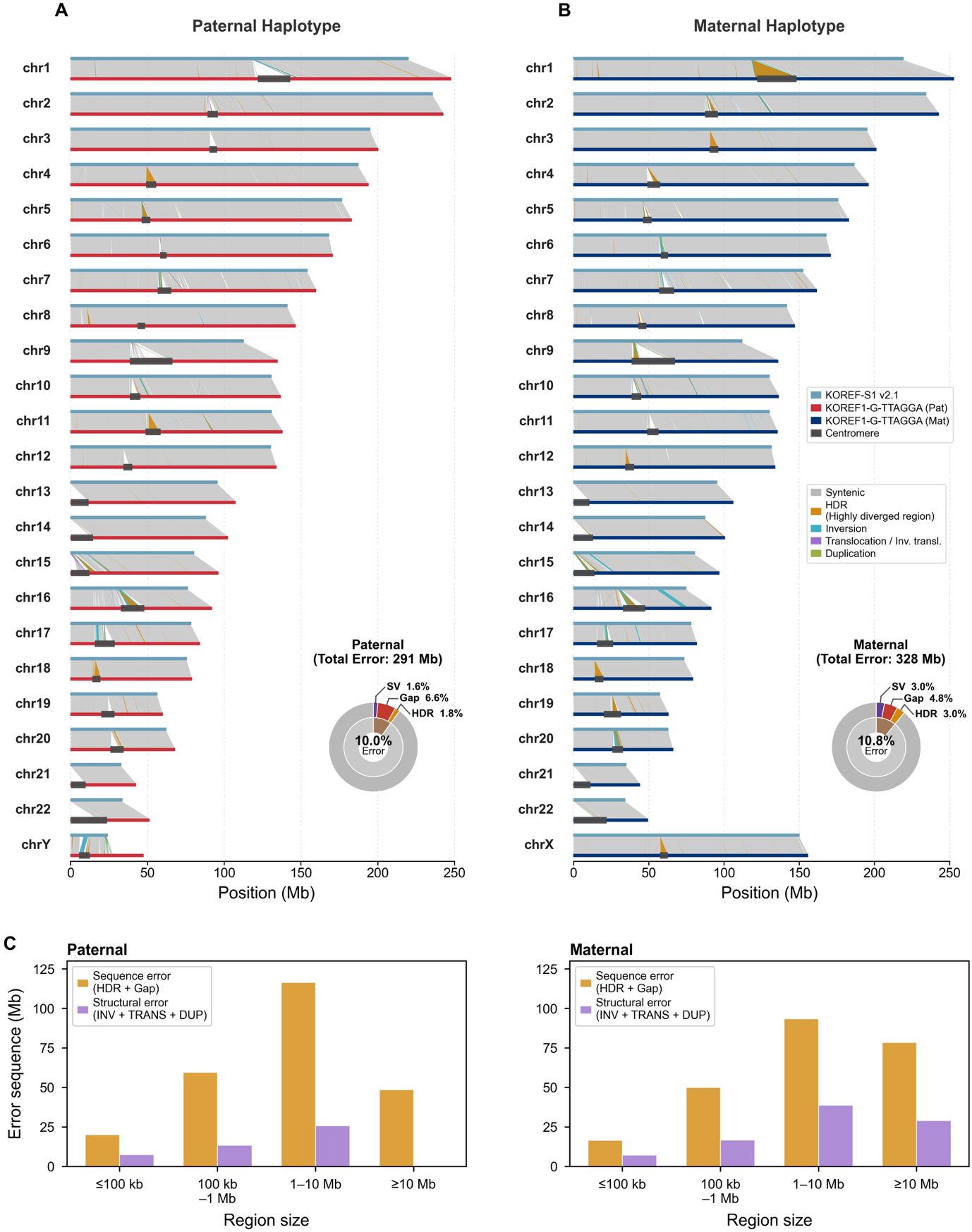

**Supplementary Figure S4. Comparison of KOREF1-G-TTAGGA with the previous KOREF assembly (KOREF-S1 v2.1).** (A, B) Whole-chromosome alignments of the previous KOREF-S1 v2.1 assembly to the (A) paternal and (B) maternal KOREF1-G-TTAGGA haplotypes. Regions are colored as syntenic, highly diverged (HDR), inversion, translocation/inverted translocation, or duplication; centromeres are indicated. Donut plots show the total sequence newly resolved in each haplotype (paternal, 291 Mb / 10.0%; maternal, 328 Mb / 10.8%), partitioned into structural variants (SV), gaps, and HDR. (C) Newly resolved error sequence (Mb) stratified by region size and by type (sequence error, HDR + Gap; structural error, INV + TRANS + DUP) for each haplotype.

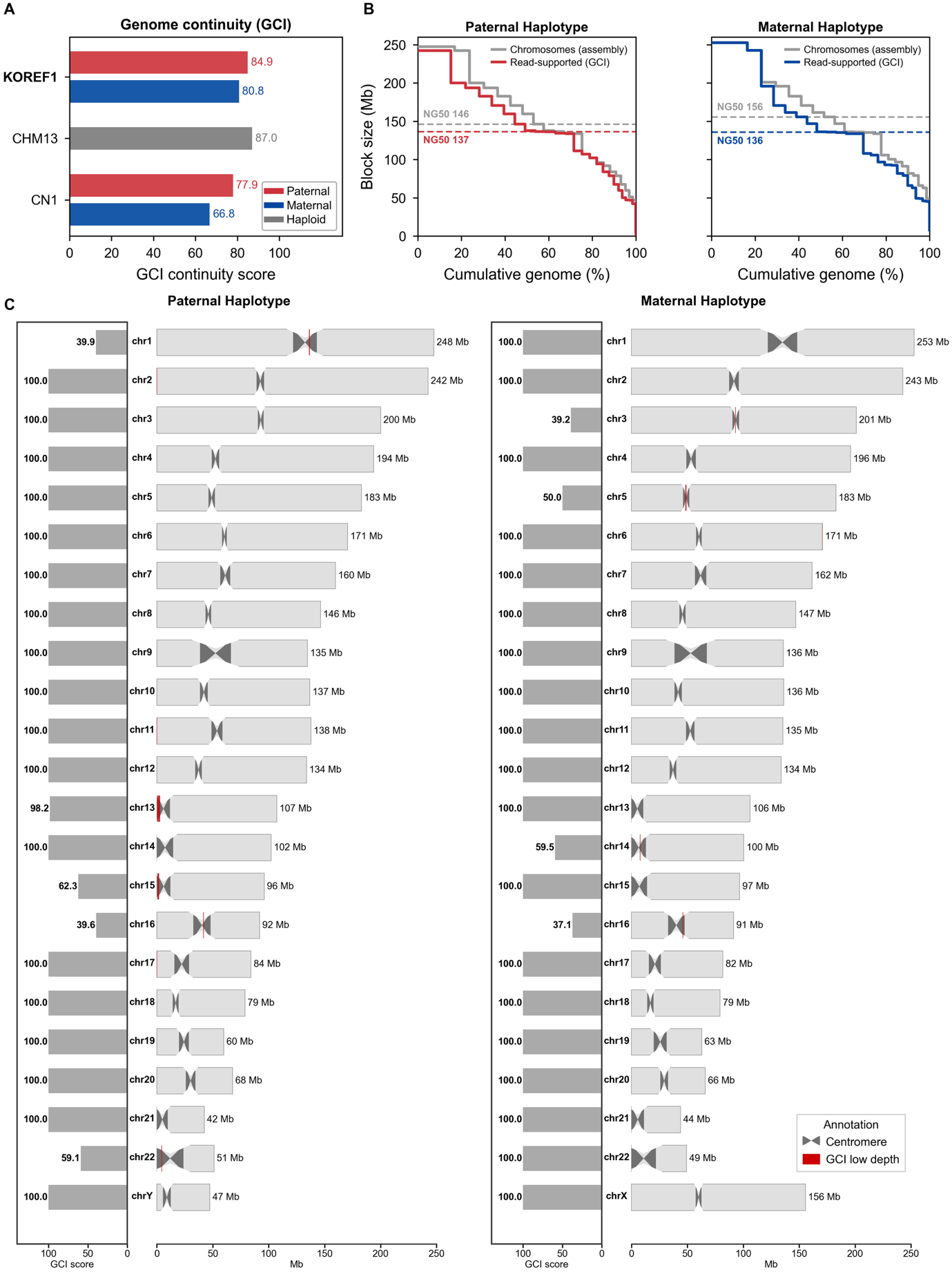

**Supplementary Figure S5. Genome continuity assessed by Genome Continuity Inspector (GCI).** (A) GCI continuity scores for KOREF1-G-TTAGGA (paternal 84.9; maternal 80.8) compared with T2T-CHM13 (87.0) and CN1 (paternal 77.9; maternal 66.8). (B) Cumulative block-size (NG50) curves for the assembled chromosomes (grey) versus read-supported blocks (GCI; colored) for each haplotype. (C) Per-chromosome GCI scores (left) and ideograms (right); centromeres and GCI low-depth regions are marked.

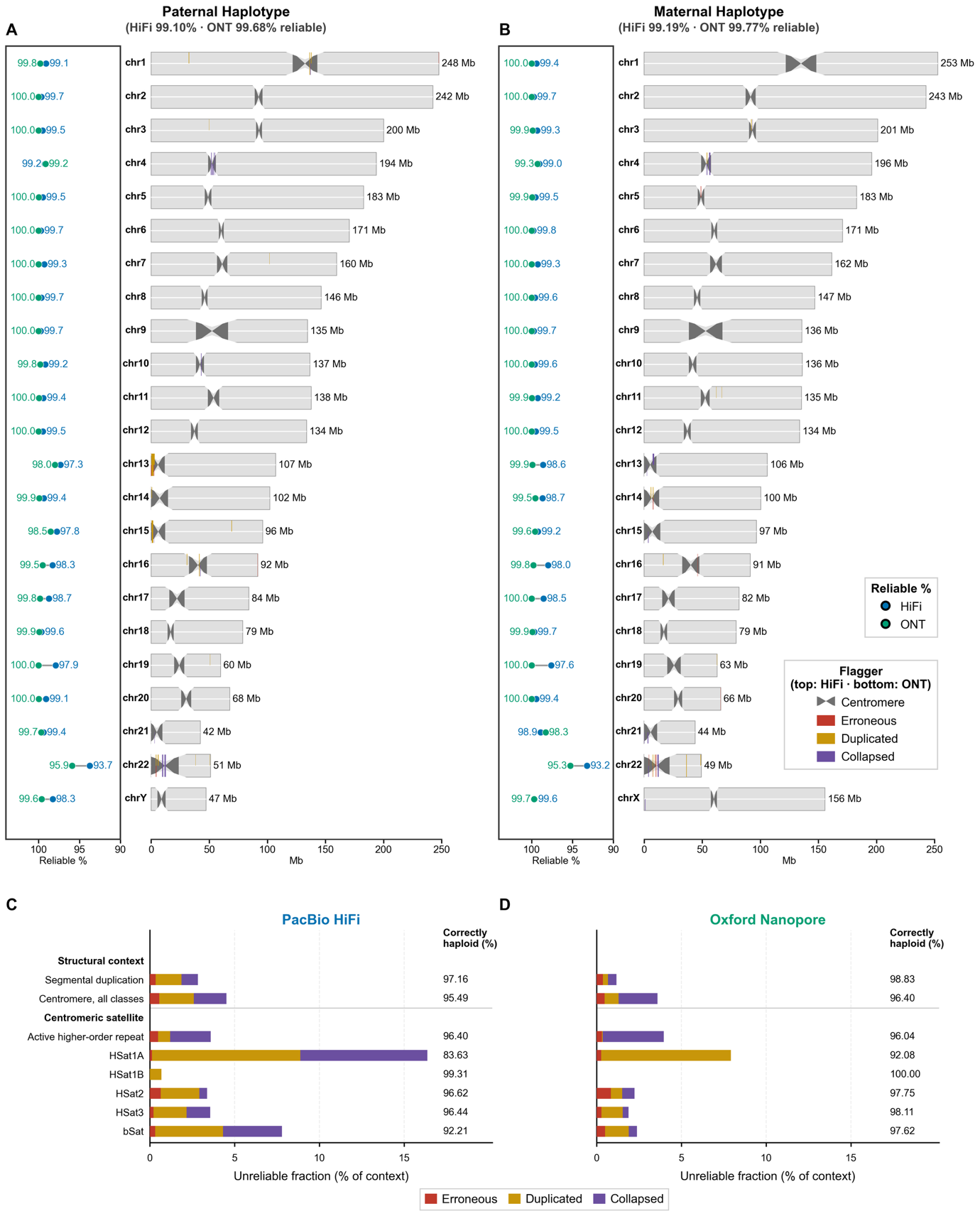

**Supplementary Figure S6. Haplotype-resolved assembly reliability from HMM-Flagger.** Per-chromosome reliable fraction (%) estimated by HMM-Flagger from independently aligned HiFi and ONT reads for the (A) paternal (HiFi 99.10%, ONT 99.68% reliable) and (B) maternal (HiFi 99.19%, ONT 99.77% reliable) haplotypes. Ideograms mark regions flagged as erroneous, duplicated, or collapsed (top, HiFi; bottom, ONT); centromeres are indicated. (C, D) Flagger read-depth reliability by sequence context, PacBio HiFi (C) and Oxford Nanopore (D). Bars give the unreliable fraction of each genomic class, so bar length is the total unreliable percentage. The correctly haploid percentage is listed at the right of each panel. Rows below the divider are the satellite classes that make up the centromere row above it. Values pool both haplotypes.

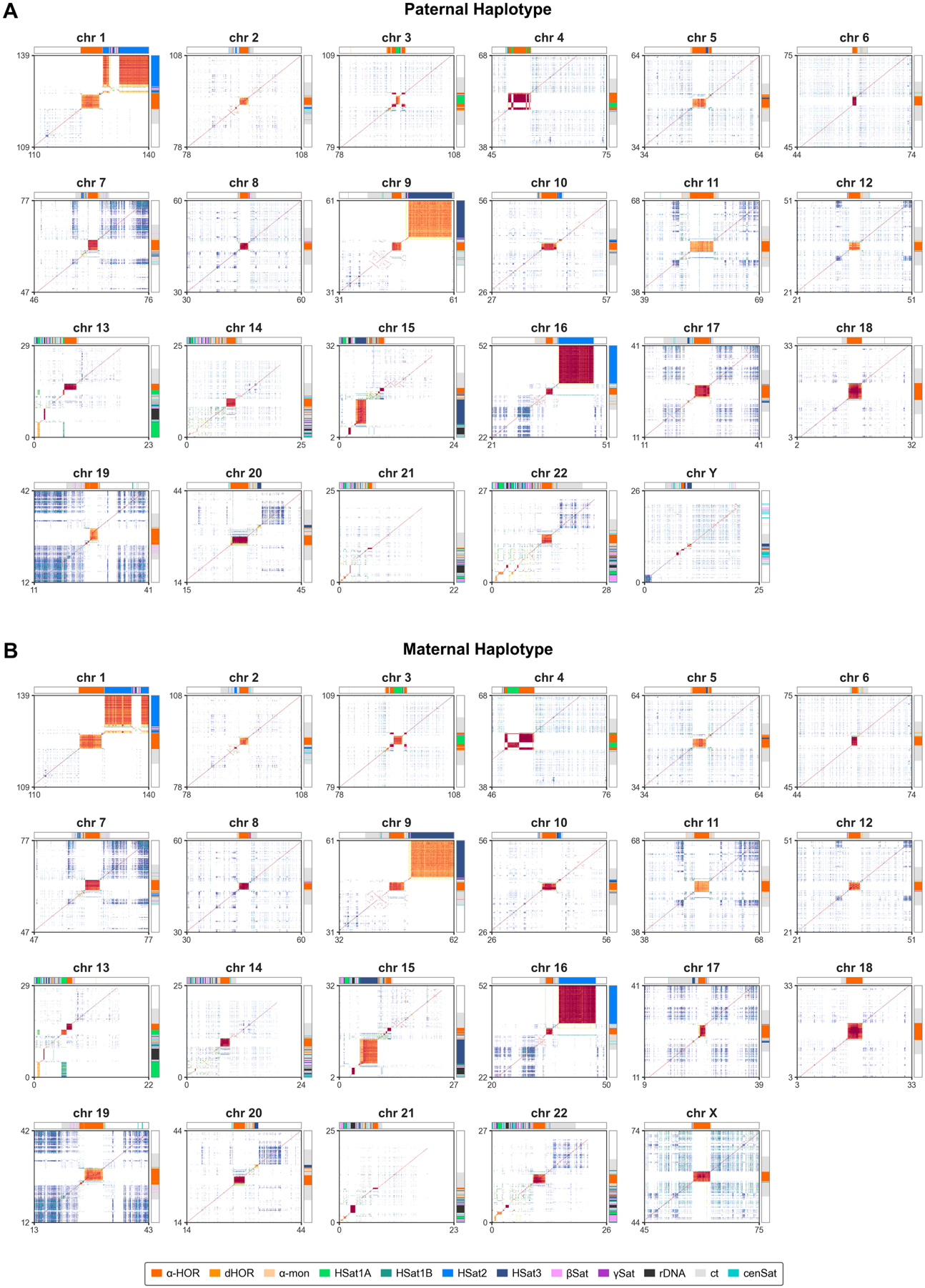

**Supplementary Figure S7. Centromere and satellite organization of KOREF1-G-TTAGGA relative to T2T-CHM13.** ModDotPlot comparison of the (peri)centromeric region of each chromosome for the (A) paternal and (B) maternal haplotypes. Each panel compares a 30 Mb window centred on the centromere, KOREF1 on the horizontal axis and CHM13 on the vertical axis, in 30 kb windows; colour gives the ModDotPlot window identity. Annotation tracks give the satellite classes, above each panel for KOREF1 and to its right for CHM13: alpha-satellite higher-order repeat (α-HOR), divergent HOR (dHOR), alpha-monomer (α-mon), human satellites (HSat1A, HSat1B, HSat2, HSat3), beta-satellite (bSat), gamma-satellite (gSat), rDNA, centromeric satellite (cenSat) and centromere transition (ct).

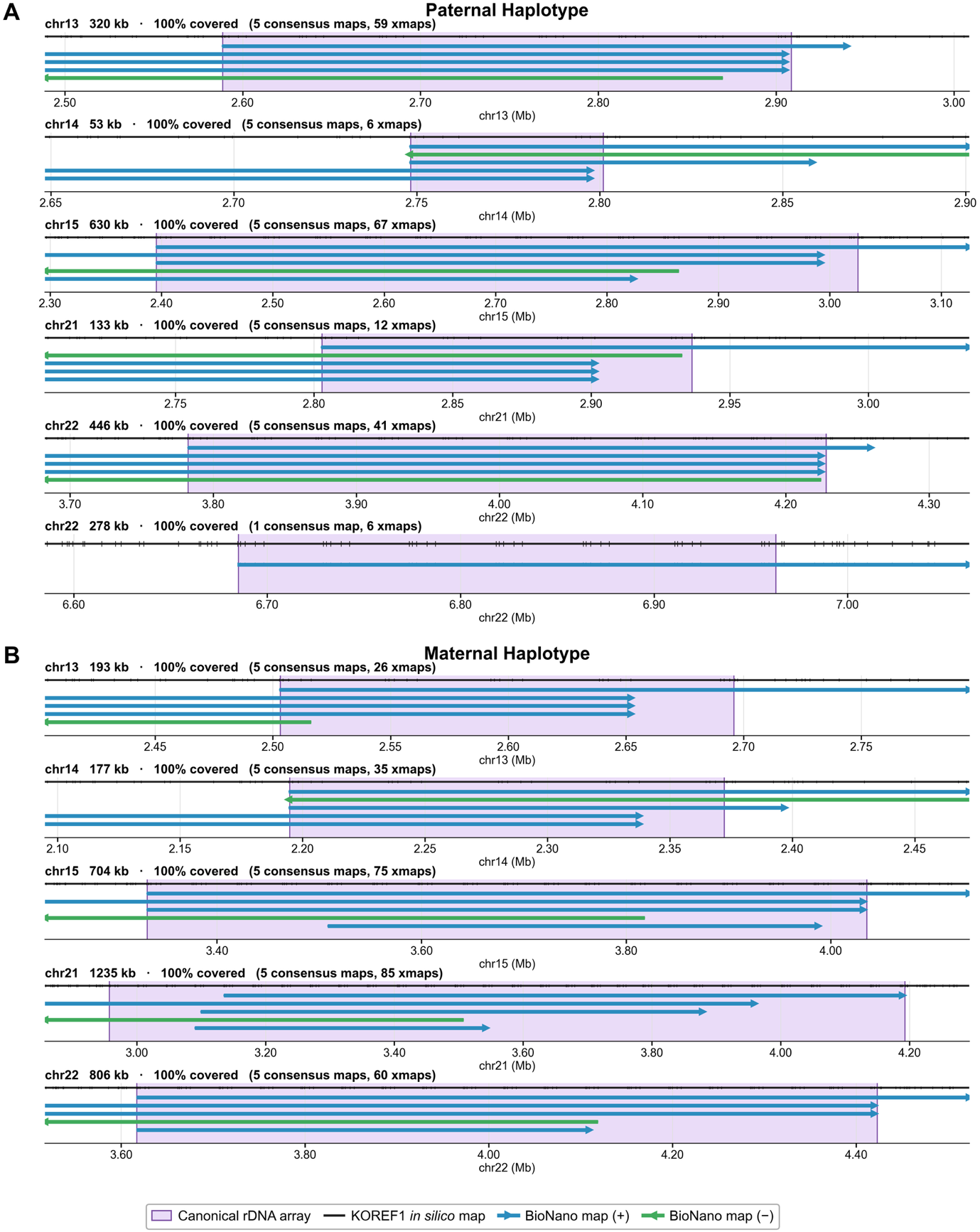

**Supplementary Figure S8. Optical-map validation of ribosomal DNA (rDNA) arrays.** Bionano optical maps spanning the canonical rDNA arrays (purple) on the acrocentric chromosomes (chr13, 14, 15, 21, 22) for the (A) paternal and (B) maternal haplotypes. The KOREF1 in silico map (black) and BioNano consensus maps in forward (+, blue) and reverse (-, green) orientation are aligned; each array is 100% covered. Array size and the number of consensus maps and molecule (xmap) alignments are indicated per panel.

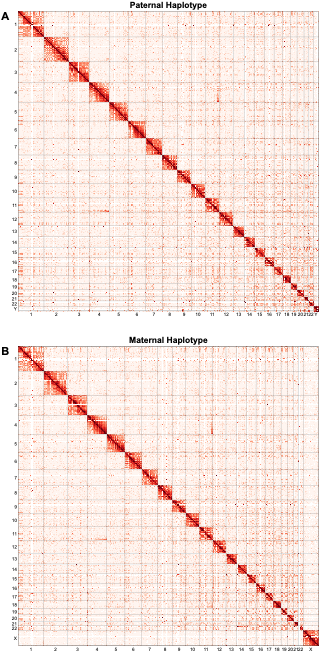

**Supplementary Figure S9. Genome-wide Hi-C contact maps.** All-chromosome Hi-C contact matrices for the (A) paternal and (B) maternal haplotypes. The block-diagonal pattern with minimal off-diagonal signal indicates chromosome-scale phasing and structural integrity.

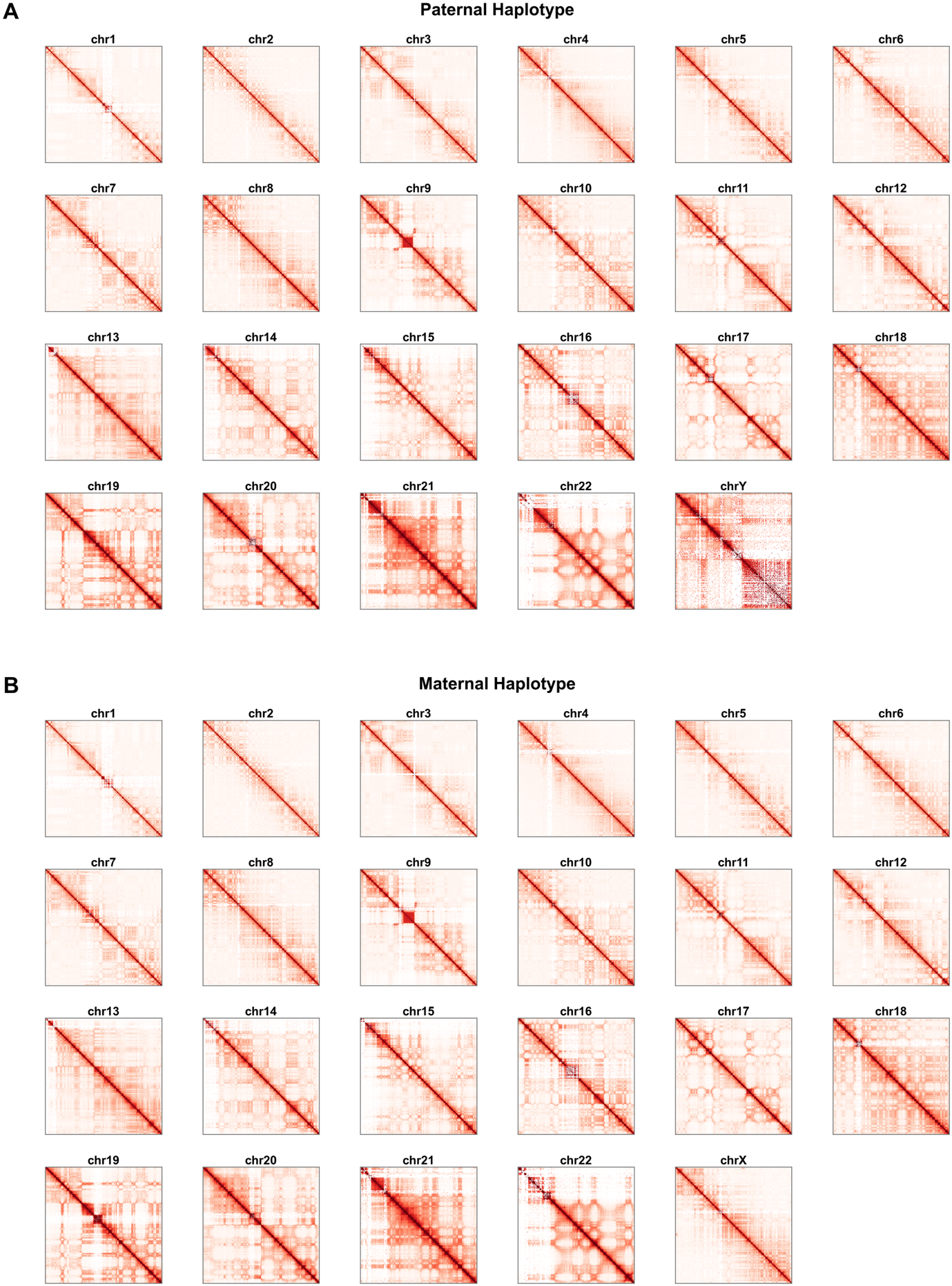

**Supplementary Figure S10. Chromosome-wise Hi-C contact maps of the paternal and maternal KOREF1-G-TTAGGA.** Hi-C interaction matrices are shown for each chromosome of the (A) paternal and (B) maternal assemblies. Contact frequencies were normalized using the Knight-Ruiz (KR) matrix balancing method and visualized on a log10 scale, where darker red indicates higher interaction frequency. Continuous diagonal patterns across all chromosomes confirm correct scaffolding and chromosomal integrity, while distinct box-like or grid-shaped patterns within certain regions correspond to repetitive or segmentally duplicated sequences, such as centromeric or satellite-rich loci.

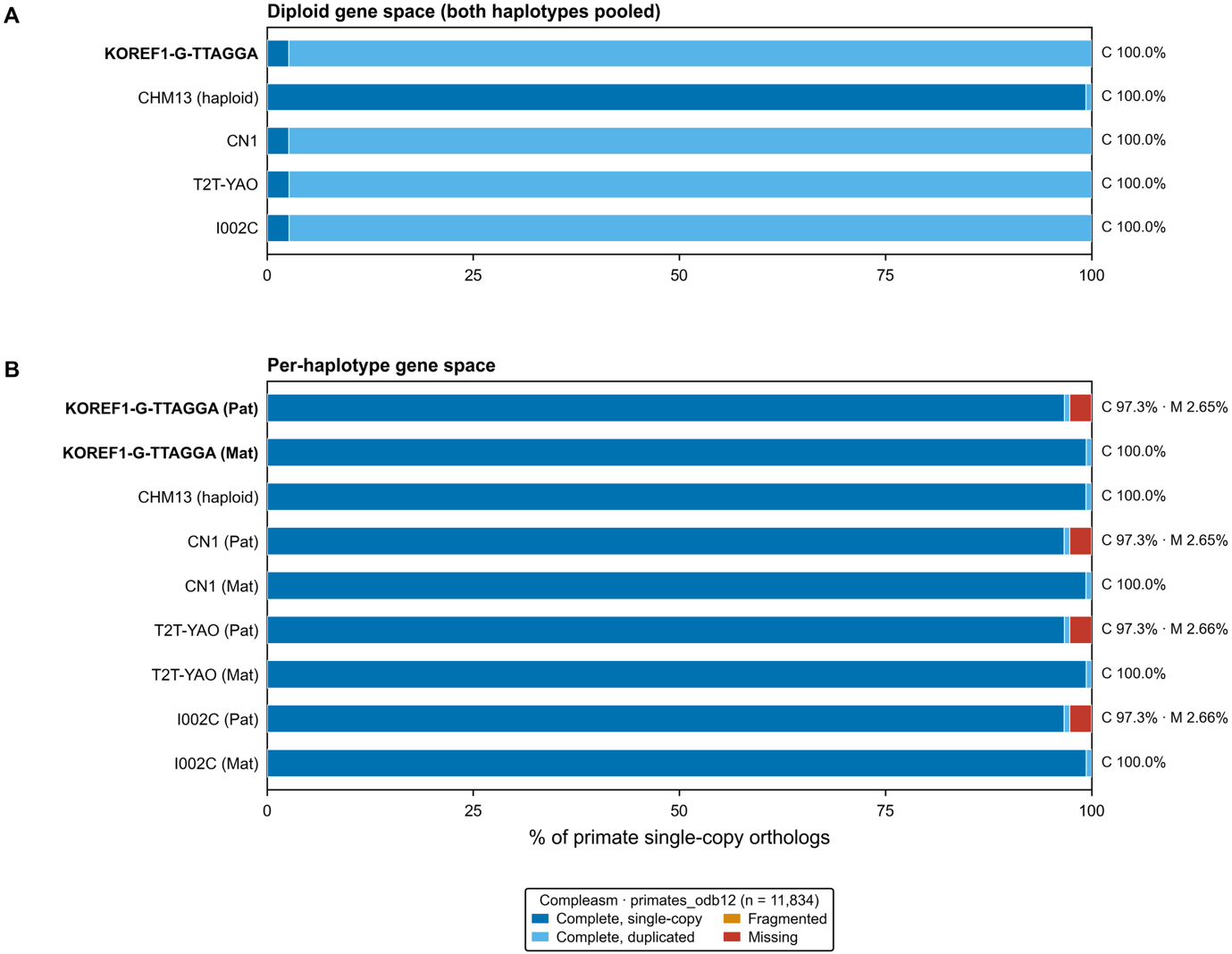

**Supplementary Figure S11. Gene completeness assessment of KOREF1-G-TTAGGA and other high-quality human genome assemblies.** Conserved single-copy ortholog completeness was evaluated using compleasm with the OrthoDB v12 primate_odb12 database (N=11,834). Bars represent the counts of complete (single-copy and duplicated), fragmented (partial and chimeric), and missing orthologs for each assembly, for (A) diploid gene space (both haplotypes pooled) and (B) per-haplotype gene space. Both paternal and maternal KOREF1-G-TTAGGA exhibit comparable completeness relative to other telomere-to-telomere human references (CHM13 v2, T2T-YAO v1.1, CN1 v1.0.1, and I002C v0.7), indicating near-complete representation of conserved primate genes.

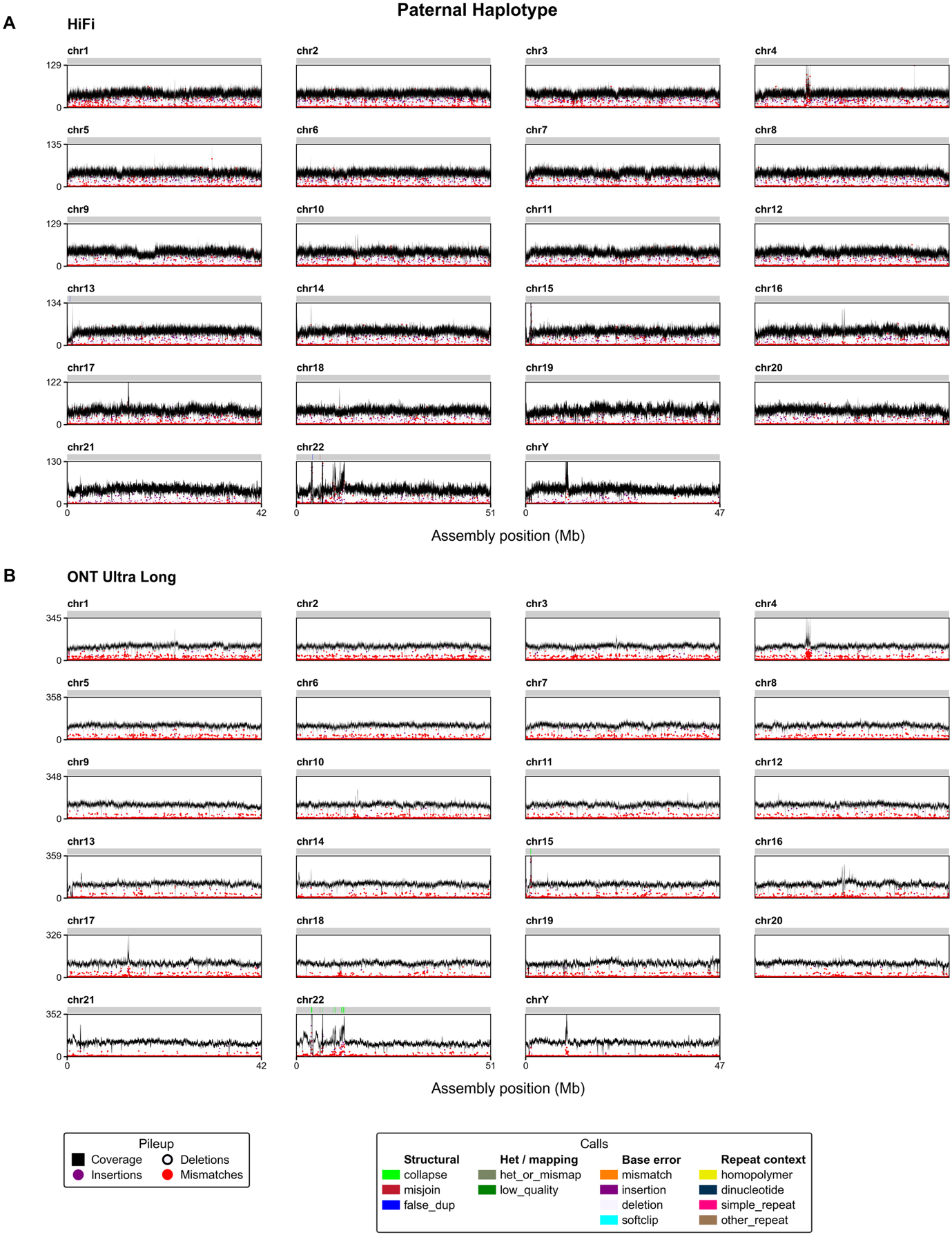

**Supplementary Figure S12. Read-pileup assembly evaluation (NucFlag) - paternal haplotype.** Coverage pileups of (A) HiFi and (B) ONT ultra-long reads aligned to the paternal haplotype for each chromosome. Track colors denote pileup features (coverage, deletions, insertions, mismatches) and called issues grouped as structural (collapse, misjoin, false duplication), heterozygous/mapping, base error, and repeat context.

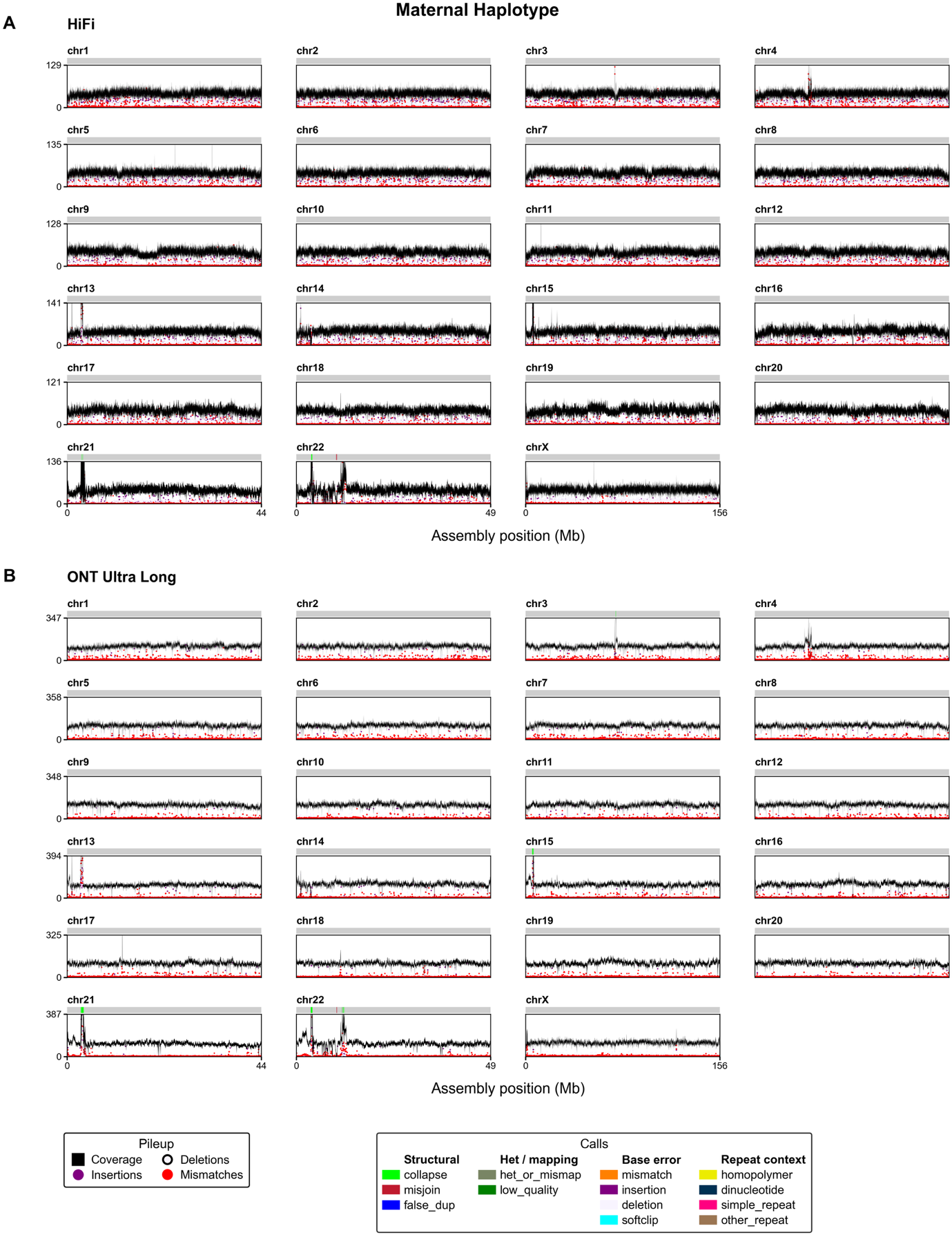

**Supplementary Figure S13. Read-pileup assembly evaluation (NucFlag) - maternal haplotype.** Coverage pileups of (A) HiFi and (B) ONT ultra-long reads aligned to the maternal haplotype for each chromosome. Track colors denote pileup features (coverage, deletions, insertions, mismatches) and called issues grouped as structural (collapse, misjoin, false duplication), heterozygous/mapping, base error, and repeat context.

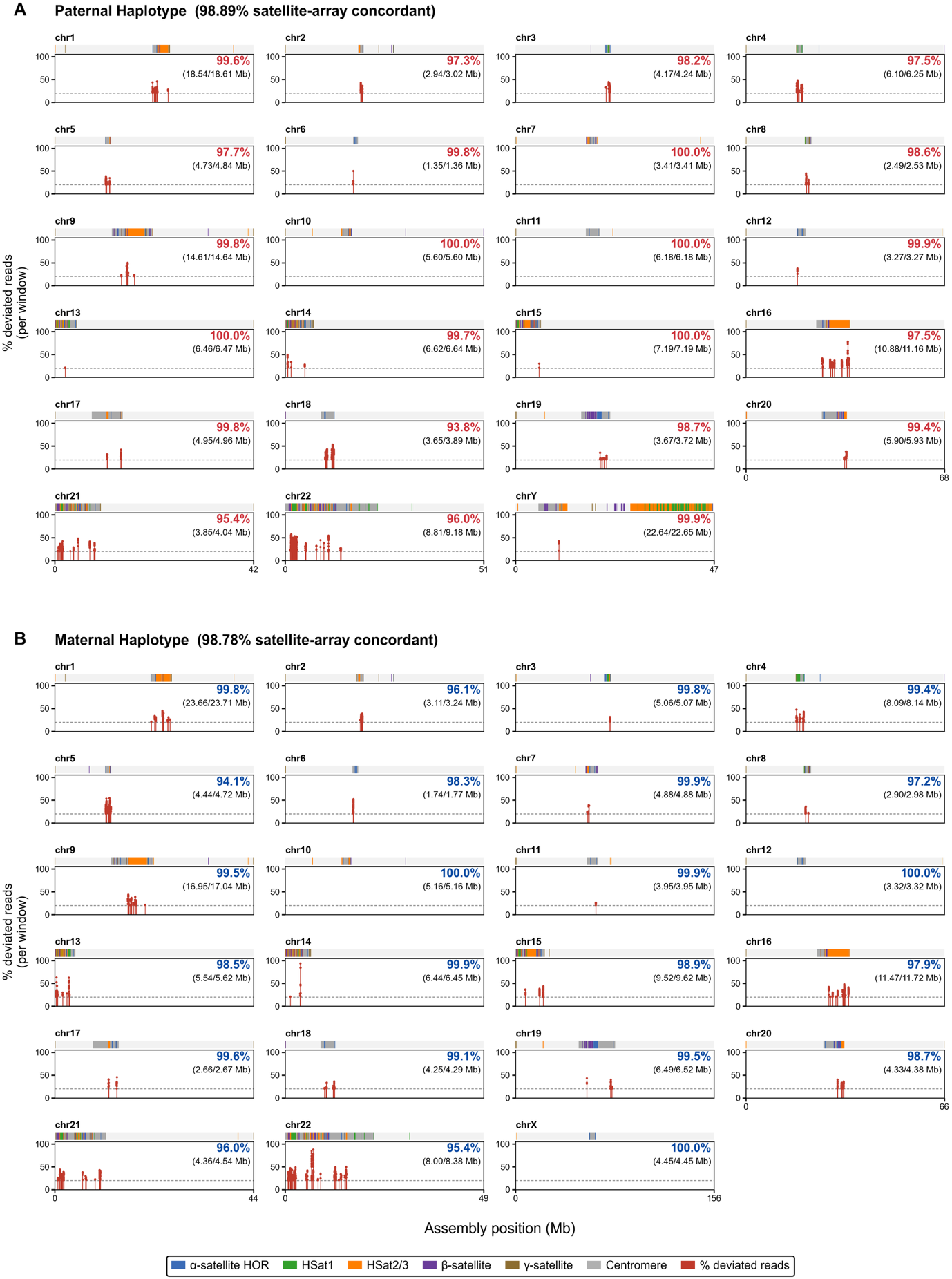

**Supplementary Figure S14. Satellite-array validation by rare k-mer mapping (VerityMap).** Percentage of deviated reads per window across satellite arrays for the (A) paternal (98.89% satellite-array concordant) and (B) maternal (98.78%) haplotypes. Annotation bars mark satellite classes (alpha-satellite HOR, HSat1, HSat2/3, beta-satellite, gamma-satellite) and centromeres; per-chromosome concordance and validated/total array size (Mb) are indicated.

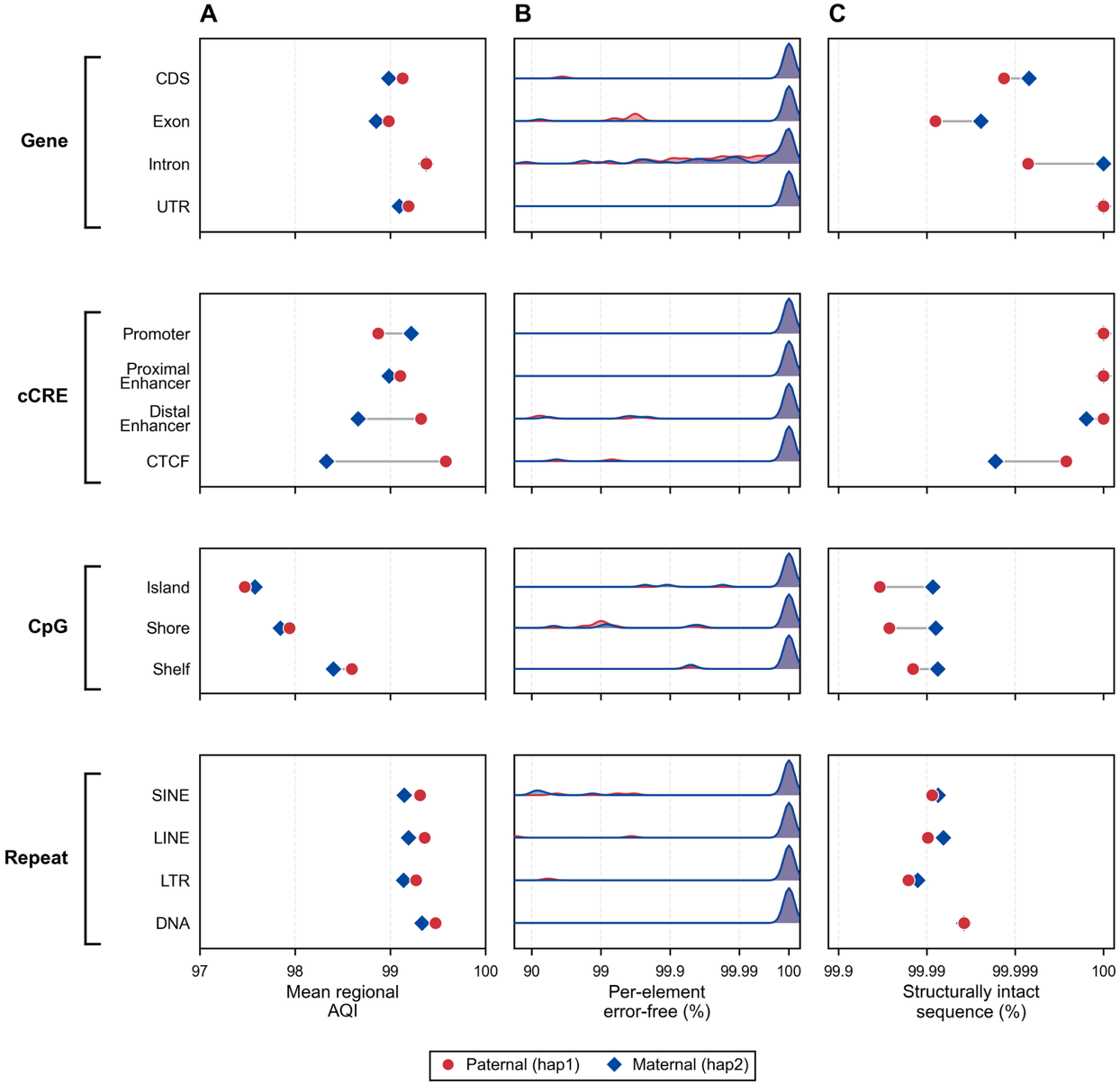

**Supplementary Figure S15. Assembly quality by functional element class (CRAQ).** CRAQ metrics for gene features (CDS, exon, intron, UTR), candidate cis-regulatory elements (promoter, proximal enhancer, distal enhancer, CTCF), CpG features (island, shore, shelf), and repeats (SINE, LINE, LTR, DNA), for the paternal (hap1, red circles) and maternal (hap2, blue diamonds) haplotypes. (A) Mean regional AQI, the CRAQ ~3 Mb window score averaged over the sequence of each class. (B) Distribution of the per-element error-free percentage (reverse-log axis, log-scaled ridge height). (C) Structurally intact sequence, the class mean of the per-element fraction free of structural error (reverse-log axis).

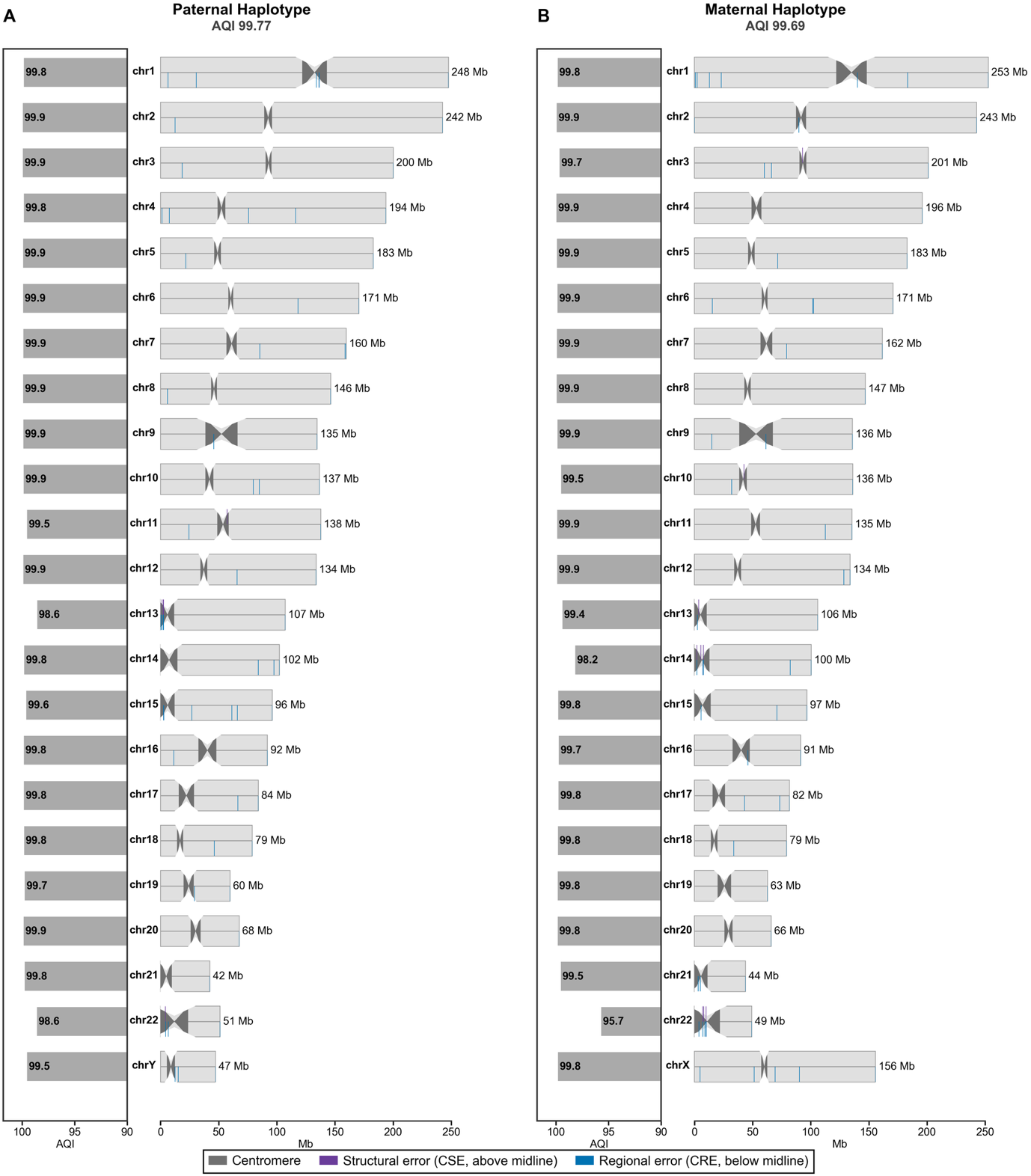

**Supplementary Figure S16. Per-chromosome assembly quality (CRAQ).** Per-chromosome CRAQ AQI for the (A) paternal and (B) maternal haplotypes. Ideograms mark structural errors (CSE, above midline) and regional errors (CRE, below midline); centromeres are indicated.
